# Generative replay for compositional visual understanding in the prefrontal-hippocampal circuit

**DOI:** 10.1101/2021.06.06.447249

**Authors:** Philipp Schwartenbeck, Alon Baram, Yunzhe Liu, Shirley Mark, Timothy Muller, Raymond Dolan, Matthew Botvinick, Zeb Kurth-Nelson, Timothy Behrens

## Abstract

Understanding the visual world is a constructive process. Whilst a frontal-hippocampal circuit is known to be essential for this task, little is known about the associated neuronal computations. Visual understanding appears superficially distinct from other known functions of this circuit, such as spatial reasoning and model-based planning, but recent models suggest deeper computational similarities. Here, using fMRI, we show that representations of a simple visual scene in these brain regions are relational and compositional – key computational properties theorised to support rapid construction of hippocampal maps. Using MEG, we show that rapid sequences of representations, akin to replay in spatial navigation and planning problems, are also engaged in visual construction. Whilst these sequences have previously been proposed as mechanisms to plan possible futures or learn from the past, here they are used to understand the present. Replay sequences form constructive hypotheses about possible scene configurations. These hypotheses play out in an optimal order for relational inference, progressing from predictable to uncertain scene elements, gradually constraining possible configurations, and converging on the correct scene configuration. Together, these results suggest a computational bridge between apparently distinct functions of hippocampal-prefrontal circuitry, and a role for generative replay in constructive inference and hypothesis testing.

## Introduction

Visual understanding is the process of inferring the relationships between objects in a visual scene to enable interaction with or reasoning about the overall configuration. It differs from visual recognition both in its computations and in the brain regions that support it. Whilst object recognition can increasingly be accounted for by feedforward models of the visual cortices (Yamins & DiCarlo, 2016), visual understanding is a constructive process (Finke & Slayton, 1988; Orbán et al., 2008) that additionally engages at least the hippocampal formation and medial prefrontal cortex (Aly et al., 2013; Córdova et al., 2019; Ruiz et al., 2020). This is perhaps most strikingly demonstrated in natural scene understanding tasks, such as imagining novel viewpoints, in which HC and mPFC are causally engaged (Hassabis et al., 2007; Hassabis & Maguire, 2009; Lee, Buckley, et al., 2005; Lee, Bussey, et al., 2005).

Despite converging evidence for the involvement of these brain regions, the neuro-computational mechanisms that underlie flexible visual understanding are largely unknown. However, recent advances in machine learning suggest progress can be made by drawing parallels to visual planning problems (Bapst et al., 2019; Hamrick et al., 2018), rooted in model-based reinforcement learning (RL, Sutton & Barto, 1998). These computational parallels are of particular interest as model-based RL also engages HC and mPFC (Behrens et al., 2018; Brunec & Momennejad, 2021; Miller et al., 2017; Stachenfeld et al., 2017; Vikbladh et al., 2019).

Like visual understanding, model-based RL requires agents to represent the relationships (transitions) between states. However, unlike RL problems studied in the laboratory, in visual planning problems it is not conceivable that subjects might enumerate the state space or learn these transitions from experience. For example, an agent trying to construct a Lego tower out of 10 individual bricks is faced with more than 3.5 million possible brick orderings. Nevertheless, humans and other animals achieve impressive performance at such construction problems even at early stages of their development (Ullman et al., 2017). It is conceivable, therefore, that studying visual understanding problems might reveal insights that are useful for understanding more general forms of planning in more realistic natural situations.

What alternatives are there if state transitions cannot be learnt from experience? One hypothesis is representations of the current state might be built by combining together pre-learnt representations of the relational and sensory properties of that state (Manns & Eichenbaum, 2006). The state (or scene) representation is composed from its parts in a process of *inference*. By factoring the problem in this way, the complexity of the problem is massively reduced (Behrens et al., 2018). The relational properties of a tower are the same, no matter the ordering of the bricks. Such relational properties also constrain the inference in important ways – a tower structure makes building blocks on top of each other more likely that building blocks besides each other. Indeed, models built on these principles account for a variety of cellular representations in the hippocampal formation, including basic representations of space (Whittington et al., 2020). Similarly, in generative models of scene understanding (Eslami et al., 2018), embeddings of visual objects in relational positions generalise across different visual scenes. Such representations enable agents to engage in flexible compositional (visual) reasoning, a hallmark of ‘combinatorial generalisation’ allowing agents to make ‘infinite use of finite means’ (Battaglia et al., 2018).

Whilst such compositional representations might allow a compact representational form, they do not provide a mechanism for inferring the appropriate configuration (and therefore representation) of the current experience. However, further consideration of known hippocampal phenomena suggests a candidate substrate for this inference. In neural replay, sequences of cellular ensembles encoding external states of the environment are (re-)activated in time-compressed form (Diba & Buzsáki, 2007a; Foster & Wilson, 2006). Critically, the external states that are activated are non-local. Replay events have been suggested as a substrate for memory consolidation, but also for evaluating plans of the future (Gupta et al., 2010; Kay et al., 2020). One possible unifying account of these apparently disparate ideas casts replay as a mechanism for learning and sampling from generative models of the world (Foster, 2017). If true, such an account suggests that replay might not only be useful for predicting the future, but also for *understanding the present*. When constrained by sensory data, such generative models are an essential part of any inferential process (Dayan et al., 1995; Friston & Buzsáki, 2016).

To test these ideas, we designed two studies to investigate the neural representations and mechanisms that enable flexible visual reasoning. We designed a task in which subjects had to flexibly (de-)construct visual silhouettes into building blocks in their relational positions (like Lego in reverse – how was this built?). After training, we presented a set of novel silhouettes to probe the generalisable neural representations underlying visual de-construction. Using functional magnetic resonance imaging (fMRI), we found compositional representations in mPFC and the anterior hippocampus reflecting the generalisable embedding of sensory building blocks in their relational configurations. Using recently developed methods to measure replay in human Magnetoencephalography (MEG) data, we show that generative sequences of hypothetical constructions are played as the subject understands the scene. These sequences resemble hypothesis tests for visual understanding. Early sequences involve all candidate building blocks (correct and incorrect) but progress towards blocks that are known to be present. Sequences that exclusively involve unknown (inferred) blocks emerge later and only involve the correct building blocks.

Together these data suggest that known representational and computational phenomena – learnt from rodent studies of spatial reasoning and reinforcement learning - may be relevant for complex human planning and inference tasks such as visual understanding.

## Results

### Subjects solve a compositional visual construction problem

In a first study, we trained 30 human subjects on the task contingencies of constructing silhouettes out of a set of building blocks for two consecutive days (Figure 1A). Subjects learned that they had nine basic building blocks available that could be combined by placing them on-top or beside each other, without worrying about the physical stability of the constructed object (similar to playing two-dimensional Lego or Tangram, see Figure 1A). Subjects were instructed that every building block could only be used once for a given silhouette and that they had to find a solution with the minimum number of building blocks, allowing precise experimental control over the correct solutions that subjects had to infer. The resultant visual construction task has considerable computational complexity. There are at least 6*10^12^ ways of connecting the nine building blocks, implying that an exhaustive state space representation including all possible configurations is computationally intractable.

**Figure 1.**
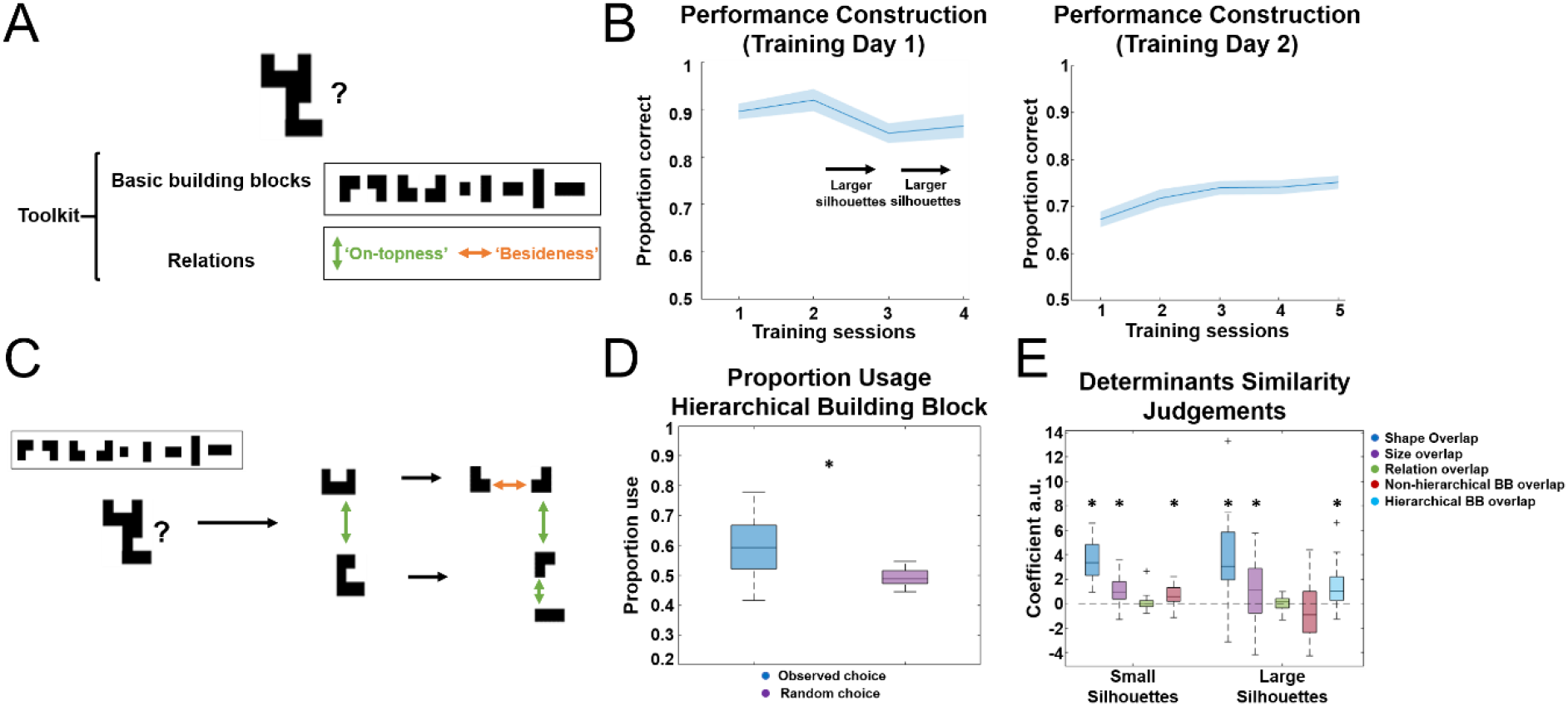
Paradigm and behavioural training (A) On two consecutive days, subjects were trained that they had nine basic building blocks available, which could be flexibly combined by placing one building block on-top (below) or beside (left or right) of another building block. (B) On the first day of training, subjects had to construct a given target shape by selecting correct building blocks and subsequently placing and connecting them in the correct way to construct a target silhouette. The complexity of the target silhouettes increased gradually, and subjects achieved overall high performance in this task. On the second day, subjects performed a more challenging task where they had to select the correct building blocks to construct a given target silhouette within a restricted time window, ensuring subjects would become fast and efficient at solving this task. Again, subjects achieved high performance and improved throughout those training sessions. (C) We included an implicit hierarchical structure in the task, such that large silhouettes could often be decomposed into hierarchical building blocks. These hierarchical building blocks were never introduced explicitly, but allowed for a more efficient construction of larger objects once learned. (D) Subjects displayed a preference for such ‘hierarchical chunking’, such that on the second training day they used such hierarchical building block configurations to construct larger silhouettes more often than predicted by chance. (E) At the end of the experiment, subjects completed a behavioural questionnaire to indicate similarity judgements between silhouettes. We found that these similarity judgements were influenced by visual similarity, namely shape (pixel) and size overlap, but also by ‘construction similarity’, namely by the overlap of (basic/hierarchical) building blocks (BB) across (small/large) silhouettes.

Nevertheless, subjects managed to solve the most basic version of the task immediately. Figure 1B shows participants’ performance on the two days of training. On day 1 (left), subjects were trained on the actual construction process. They were presented with target silhouettes that increased in complexity across training sessions, and subsequently had to select adequate building blocks and construct the target silhouettes by moving and connecting the building blocks in the correct way. Already in the first training session subjects achieved high overall performance when constructing (small) silhouettes (mean proportion correct: 0.89, std: 0.09), which remained stable in the subsequent sessions where the size of the target silhouettes increased gradually. On day 2 (right), subjects received a modified training task that ensured they became quick and efficient at this construction process. In this task, subjects were presented with a target silhouette and had to select the correct building blocks to construct this silhouette within a short amount of time, without having to actually construct the target silhouette. Again, subjects displayed high performance that gradually increased over time. This motivates the question about the underlying neural code that allows subjects to efficiently solve this task and generalise their knowledge across different construction problems.

To test whether generalisable neural representations extend across different hierarchical levels, we added an additional layer to the task. With ongoing experience, subjects could learn that larger silhouettes can often be decomposed into smaller recurring visual chunks, which are themselves built using two basic building blocks (Figure 1C bottom). Thus, subjects were implicitly exposed to a set of ‘hierarchical’ building blocks, which facilitated an efficient decomposition of larger visual silhouettes. Analysis of participants’ behaviour on the second day of training provides evidence that they indeed chose ‘hierarchical’ building blocks more often than predicted by chance when constructing larger silhouettes (Figure 1D right, see also Supplementary Figure 1).

We also probed how subjects perceived and processed these visual silhouettes in a behavioural questionnaire at the end of the experiment (end of day 3 after the fMRI scan). To do so, we asked subjects to judge the similarity of two silhouettes to a target silhouette according to how they would *construct* this silhouette (‘do you think the silhouette on the left or on the right of the screen is more similar to the silhouette in the middle of the screen based on how you would construct them?’), using silhouettes that were used during the fMRI experiment (see Figure 2B). Using a logistic regression, we probed the effects of different silhouette features on these similarity estimates, particularly visual (shape and size overlap) and construction (overlap of basic building blocks, how they relate to each other, and overlap of hierarchical building blocks if applicable) features. Investigation of the resultant regression weights indicated that those similarity judgements were guided by basic visual similarity, namely the shape (pixel) and size overlap of the candidate silhouettes with the target silhouette. Importantly, however, we also found that the overlap of relevant building blocks accurately predicted similarity judgements. In small silhouettes that where compounds of two basic building blocks, the overlap of basic building blocks predicted subjects’ similarity judgements, whereas in large silhouettes, the overlap of hierarchical building blocks was predictive of those judgements (Figure 1E).

**Figure 2.**
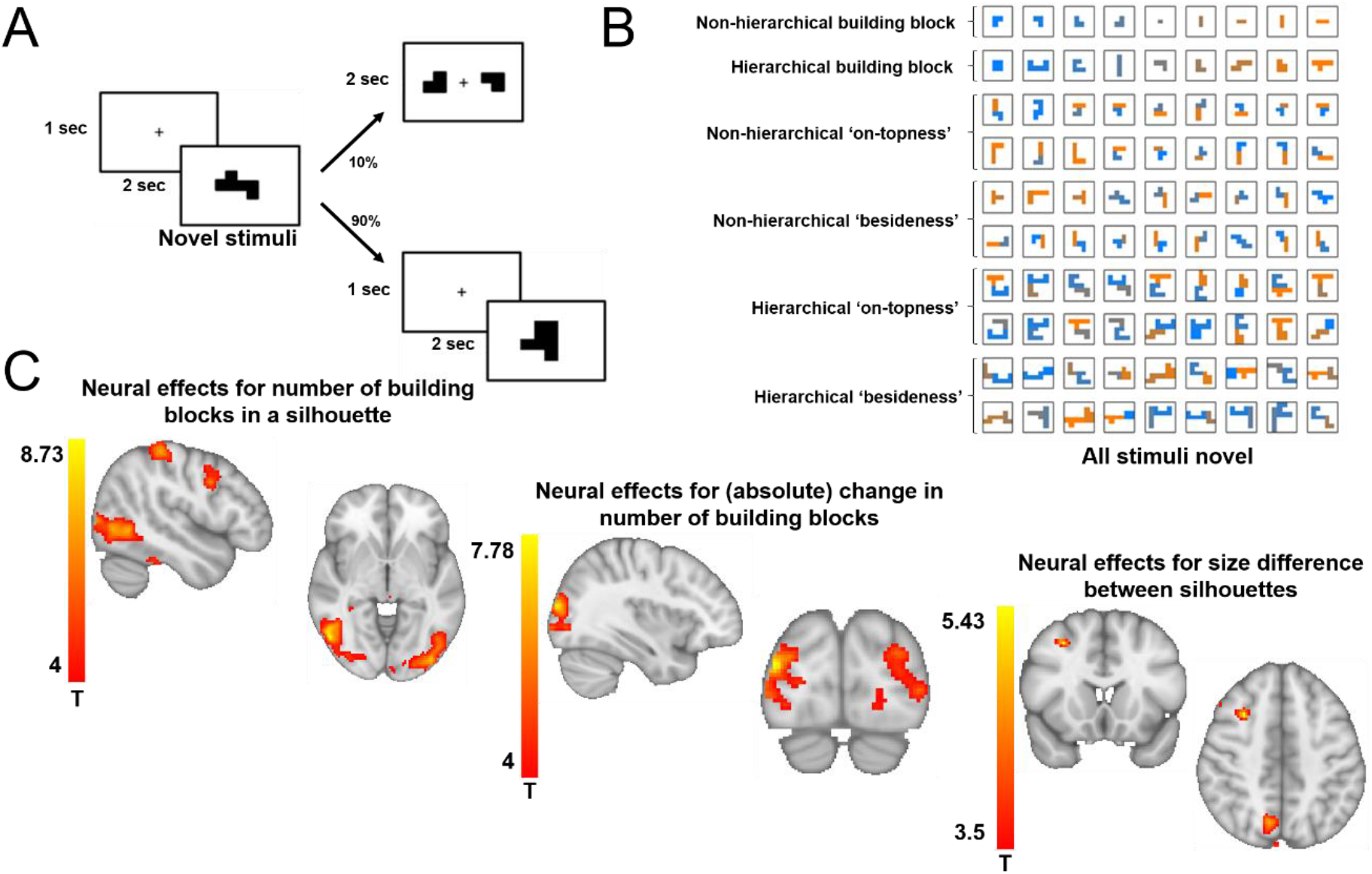
(A) After two days of training, we probed the neural representations that underlie flexible visual construction using a set of novel and previously unseen silhouettes. In the fMRI-scanner, subjects saw a silhouette for a short period of time and were instructed to infer a plan for the construction of that silhouette. To ensure that subjects engaged in this task, 10 percent of trials were followed by a catch trial, in which subjects were presented with a set of basic building blocks and had to indicate whether these blocks were part of the construction of the previous silhouette. (B) In the scanner, subjects received basic building blocks (first row), hierarchical building blocks (second row), or a novel and previously unseen compounds as construction trials. The novel compound silhouettes were either built with two basic building blocks on top of each other (third and fourth row) or beside of each other (fifth and sixth row) or with two hierarchical building blocks on top (seventh and eighth) or beside (ninth and tenth) each other. (C) We found that activity in lateral occipital, superior parietal and precentral gyrus covaried with the number of elements in a compound, providing an approximation to construction difficulty (left). We also found effects for (absolute) changes in the number of elements between consecutive silhouettes in lateral occipital cortex (middle). We did not detect any significant effects for differences in visual shape (pixels), but effects in superior parietal and frontal cortex reflecting differences in size between the individual silhouettes.

Taken together, these behavioural data suggest that subjects quickly achieved a successful representation of the generalisable task structure during behavioural training, despite the considerable computational complexity of the task.

### Visual planning signals underlying the flexible construction of silhouettes

After two sessions of training on separate days, we measured the neural representations underlying the flexible construction of silhouettes using fMRI. In the scanner, subjects saw silhouettes for a short period of time and were instructed to infer a plan to construct these silhouettes. To ensure that subjects actively engaged in the mental construction, 10% of all trials were followed by a screen probing the construction of the previous silhouette (see Figure 2A), and subjects were paid in accordance with their performance in these questions. Despite the challenging nature of the task and the short time period for the construction and probe trials, subjects achieved above chance accuracy in these probe trials (mean reaction time: 1305ms, proportion correct: 0.65). We selected silhouettes whose construction features (particular building blocks in particular relational positions) and visual features, (such as the size or visual shape of the silhouette) were de-correlated (see Supplementary Figure 2). In the scanner, trials included (basic and hierarchical) building blocks as well as compound silhouettes. Critically, these compounds were novel silhouettes that had never been experienced during training. Those compounds could either be small, i.e., built with two basic building blocks, or large, i.e., built with two hierarchical building blocks, where the underlying building blocks could either be connected via ‘on-topness’ or ‘besideness’ (Figure 2B).

Initially, we probed for effects of basic (visual) processing during the mental construction of a silhouette (Figure 2C). We found strong effects for activity in lateral occipital cortex (peak MNI [52 - 66 -4], *t*_*peak*_= 8.73), superior parietal cortex (BA7, peak MNI [-22 -74 58], *t*_*peak*_= 8.04), and precentral gyrus (peak MNI [48 6 34], *t*_*peak*_= 6.37, peak MNI [-48 -2 34], *t*_*peak*_= 5.08) that covaried with the number of basic building blocks in a given silhouette, serving as an approximation to task difficulty and engagement in the construction process (left). We also observed strong effects in lateral occipital cortex for (absolute) changes in the number of building blocks between consecutive silhouettes (peak MNI [-26 -94 14], *t*_*peak*_= 6.08, peak MNI [38 -88 16], *t*_*peak*_= 7.78). In this and all following imaging analyses, we controlled for shape (pixel) and size overlap effects as potential visual confounds in our analyses. We did not detect any significant effects for differences in pixel overlap between visual silhouettes, but found effects for size differences in superior parietal cortex (peak MNI [10 -68 46], *t*_*peak*_= 4.74), and medial frontal gyrus (peak MNI [30 16 46], *t*_*peak*_= 4.74). All effects are cluster-corrected at p<0.001. This suggests that the component building blocks are reflected in visual activity over and above the basic visual properties of the silhouette.

### Compositional neural representations in the medial prefrontal cortex and hippocampal formation

Importantly, our task design allowed us to go further than probing the effects of visual processing, and investigate neural representations that facilitate the internal construction of the object from its component parts – an example of compositional reasoning (Lake et al., 2015; Lake, 2019). Specifically, our key hypothesis concerned the neural representations of building blocks in specific relational configurations that can be generalised across different stimuli, such as knowing what it means for an object to be on top of other objects. Such a representation implies neural patterns that encode specific *conjunctions* of building blocks in a given relational position, for example a building block on-top of but not below of another building block. Such conjunctive representations can be flexibly combined, such as combining ^*<u>W</u>*^ (W on top of something) with 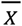 (X below of something), together providing a neural code for the composed object 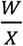 (note that these are spatial relations of blocks, not fractions).

This hypothesis predicts that neural representations for specific building blocks in a specific relational position should generalise across different silhouettes. Critically, this allows us to predict specific relational configurations of building blocks using representations of other configurations in a ‘silhouette algebra’ (Eslami et al., 2018), as illustrated in Figure 3A (see Supplementary Figure 3 for all trials). For example, given building blocks WXYZ, silhouette algebra says 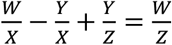. Notably, we can perfectly control for the building blocks that are used by asking that the left-hand side of the equation predicts 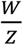 (target) but not 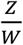 (reference) that uses the same blocks. Such a representation is *compositional* - it uses the same representations to encode the blocks in different constructed silhouettes – but also *conjunctive* as these representations differ depending on the relational position of the blocks.

**Figure 3.**
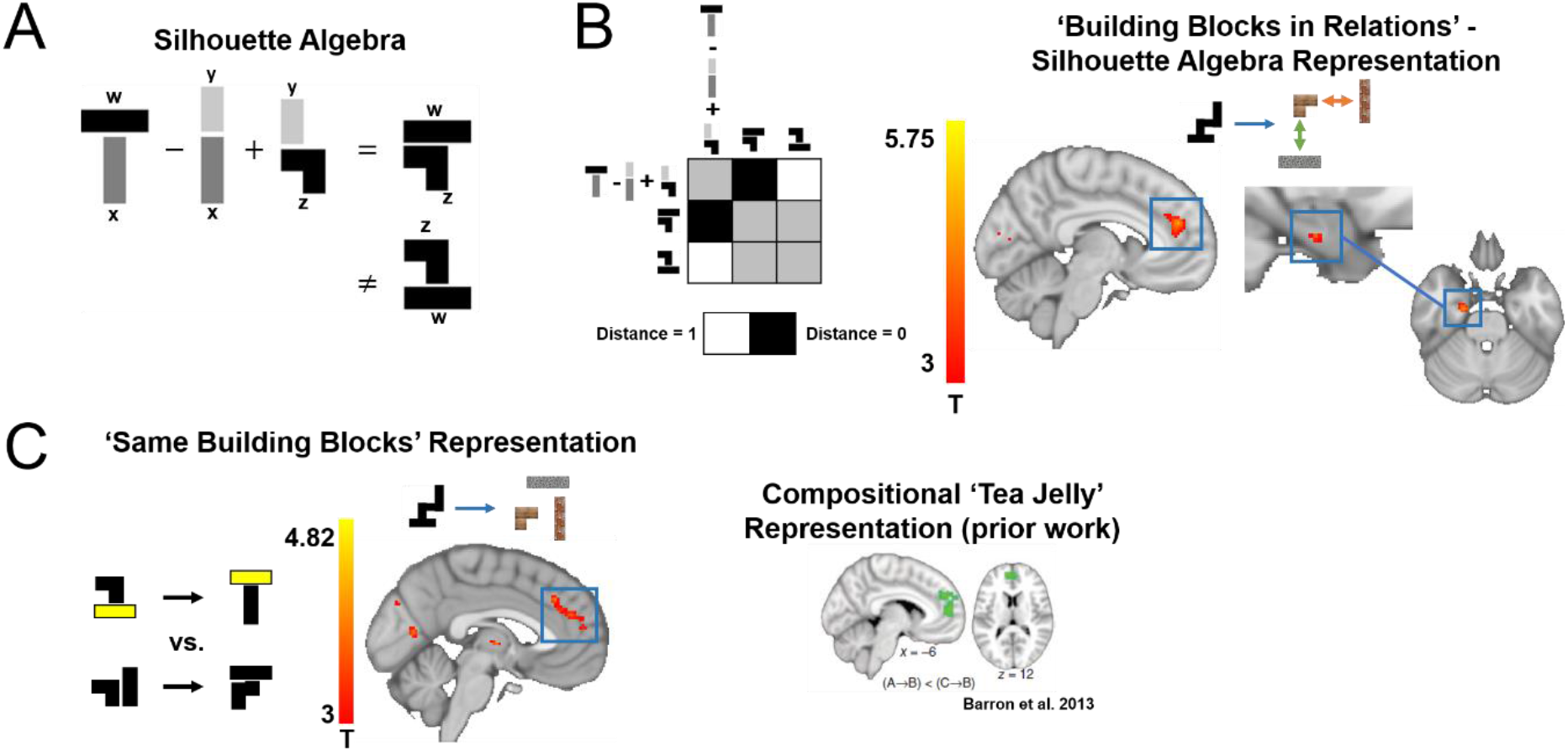
(A) ‘Silhouette algebra’ (cf., Eslami et al., 2018) to test for a conjunctive output representation in fMRI. We designed an analysis to test for generalisable representations of individual building blocks in specific relational positions by performing algebraic operations with neural representations for different silhouettes. For given building blocks WXYZ, the silhouette algebra predicts that 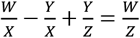 (note that these are spatial relations of blocks, not fractions). Specifically, when subtracting the neural representation of a silhouette with building block Y on top of building block X from a silhouette with building block W on top of building block X, then building block X cancels out (dark grey bricks) leaving (building block W on top) – (building block Y on top). If we now add the representation for a silhouette with building block Y on top of building block Z, then building block Y (light grey) cancels out leaving (building block W on top) + (building block Z below). Under a conjunctive representation, this algebraic term should be predictive of the actual silhouette with building block W on top of building block Z (target) but not a silhouette with building block Z on top of building block W (reference). (B) Left: We tested in which brain regions such algebraic terms are predictive of target silhouettes but not reference silhouettes using RSA, where we assessed whether the distance (defined as 1-correlation between activity patterns) between algebraic terms and target silhouettes is smaller than between algebraic terms and reference silhouettes (see methods for details). Right: We found significant effects in mPFC and the anterior hippocampus, extending into entorhinal cortex, suggestive of a conjunctive representation of building blocks in specific relational positions. (C) Using repetition suppression, we probed the neural representations encoding for individual building blocks in a given construction problem as reported in previous work when subjects had to imagine and evaluate novel food items (Barron et al., 2013). In regions encoding such representations, we expect higher suppression for transitions between silhouettes that share building blocks compared to transitions of silhouettes that use different building blocks. We found the strongest suppression effects in medial prefrontal cortex (red), highly overlapping with representations underlying the construction and evaluation of novel food items reported earlier (green, Barron et al., 2013, Figure 2C).

We used searchlight representational similarity analysis (RSA) (Kriegeskorte et al., 2008) to assess whether we can predict relational configurations in many such equations (see Supplementary Figure 3). In each searchlight, we computed the activity pattern of the terms left-hand side (‘silhouette algebra’) of the equation by combining the different silhouettes according to the signs in the equation (Figure 3A). We then computed the representational distance between this predicted pattern and the measured patterns for the target and reference silhouettes. We then compared this to a template distance matrix that asked whether the ‘silhouette algebra’ representation was more similar to the target than the reference (Figure 3B), whilst controlling for visual (shape and size) similarity.

Across the two hierarchical levels, this ‘silhouette algebra’ analysis predicted voxelwise patterns in the mPFC (peak MNI [41 78 46], *t*_*peak*_= 4.67, p = 0.045 based on a cluster-mass FWE-corrected whole brain nonparametric permutation test). We found that these effects are stronger for the non-hierarchical silhouettes alone in the mPFC (peak MNI [40 76 42], *t*_*peak*_= 5.35, *p* = 0.022 based on a cluster-mass FWE-corrected whole brain nonparametric permutation test) and also in anterior hippocampus, extending into entorhinal cortex (peak MNI 29 51 21, *t*_*peak*_ = 4.69, *p* = 0.028 based on voxel-wise FWE-corrected nonparametric permutation test corrected for the bilateral hippocampal formation), as shown in Figure 3B (right). The latter finding is closely aligned with recent evidence that translations in abstract stimulus space can predict representations of stimuli in the anterior hippocampus, highly overlapping with our effects (Morton et al., 2020). We did not detect significant effects for a hierarchical ‘silhouette algebra’ alone.

This suggests that mPFC and anterior HC have representations that reflect the building blocks in their correct configuration. Previous work has also highlighted representations in mPFC and hippocampus when constructing novel items (Barron et al., 2013; Bongioanni et al., 2021). Specifically, this work has found a critical involvement of the mPFC and hippocampus in evaluating novel food items, such as ‘tea jelly’ built out of ‘tea’ and ‘jelly’. A central difference to the algebra analysis reported above it that a ‘tea jelly’ neural code does not differentiate between different relational embeddings of the individual building blocks that were used to construct a specific food item.

We tested for an analogous ‘tea jelly’ representation in our visual construction task by disregarding the individual relational positions of individual building blocks in a compound, and simply using the overlap of individual building blocks across silhouettes as a measure of similarity instead. This can be thought of as the ‘input’ to a given construction problem, reflective of the relevant building blocks used in a given construction problem. To ensure consistency with earlier approaches, we employed cross-stimulus fMRI adaption (Barron et al., 2016; Buckner et al., 1998). Here, compositional ‘tea jelly’ representations predict stronger suppression effects for silhouettes (i.e., transitions between silhouette-trials) that share the same compared to different building blocks. Consequently, observing such effects would indicate the encoding of silhouettes according to their input building blocks.

Previous reports of such construction effects have been based on valuation tasks (Barron et al., 2013; Bongioanni et al., 2021). By contrast, our task was a visual construction paradigm with no valuation component. Despite these differences, we found repetition suppression effects for these ‘input’ representations in overlapping neural structures, particularly in the mPFC (Figure 3C, red: compositional representations underlying visual construction, green: effects from Barron et al., 2013 Figure 2C; peak MNI coordinates voxel-wise FWE-corrected and masked by effects of Barron et al., 2013: [2 52 16], *t*_*peak*_ = 3.91, *p* = 0.037).

### Temporal characteristics of compositional visual construction

Our fMRI data support the view that visual understanding problems engage the hippocampal-prefrontal circuitry. They further show that, in these brain regions, representations for visual construction share similarities with those in planning and evaluation (Barron et al., 2013; Bongioanni et al., 2021) and in spatial reasoning (Manns & Eichenbaum, 2006). This opens up the possibility that mechanisms known to represent possible *futures* in these planning contexts might also underlie hypothesis testing about possible *presents*.

One such mechanism is replay (Diba & Buzsáki, 2007a; Foster & Wilson, 2006). In rodents solving spatial tasks, hippocampal cells signal the current location of the animal, but during rest (Ambrose et al., 2016) and planning (Johnson & Redish, 2007; Pfeiffer & Foster, 2013) transiently signal sequences of remote locations. It is suggested that at least some of these events signal a roll-out of a model of the world to predict possible futures and enable choices (Gupta et al., 2010; Kay et al., 2020). Recently, we and others have developed tools to measure such sequences noninvasively in humans using MEG (Kurth-Nelson et al., 2016; Liu et al., 2019), and shown that they share many properties with rodent replay. We therefore designed an MEG experiment to probe the temporal dynamics and potential mechanisms underlying compositional visual understanding.

20 human subjects were pre-trained on a visual construction task for two consecutive days (Figure 4A). This task was similar to the task used in the fMRI above, but with two key differences. First, to optimise MEG decoding, we only had 4 building blocks, and gave each building block a unique texture. This meant we could not have a hierarchical version of the task, which would require more than 4 blocks. Second, one of the 4 building blocks was present in every silhouette (‘stable’). This was included to introduce asymmetry into possible plans, which allowed us to define the directionality of replay akin to forward and backward sequences (Liu et al., 2019, 2021). Here, these different directions translate into replay starting from the stable or present building blocks (explained in the replay section below).

**Figure 4.**
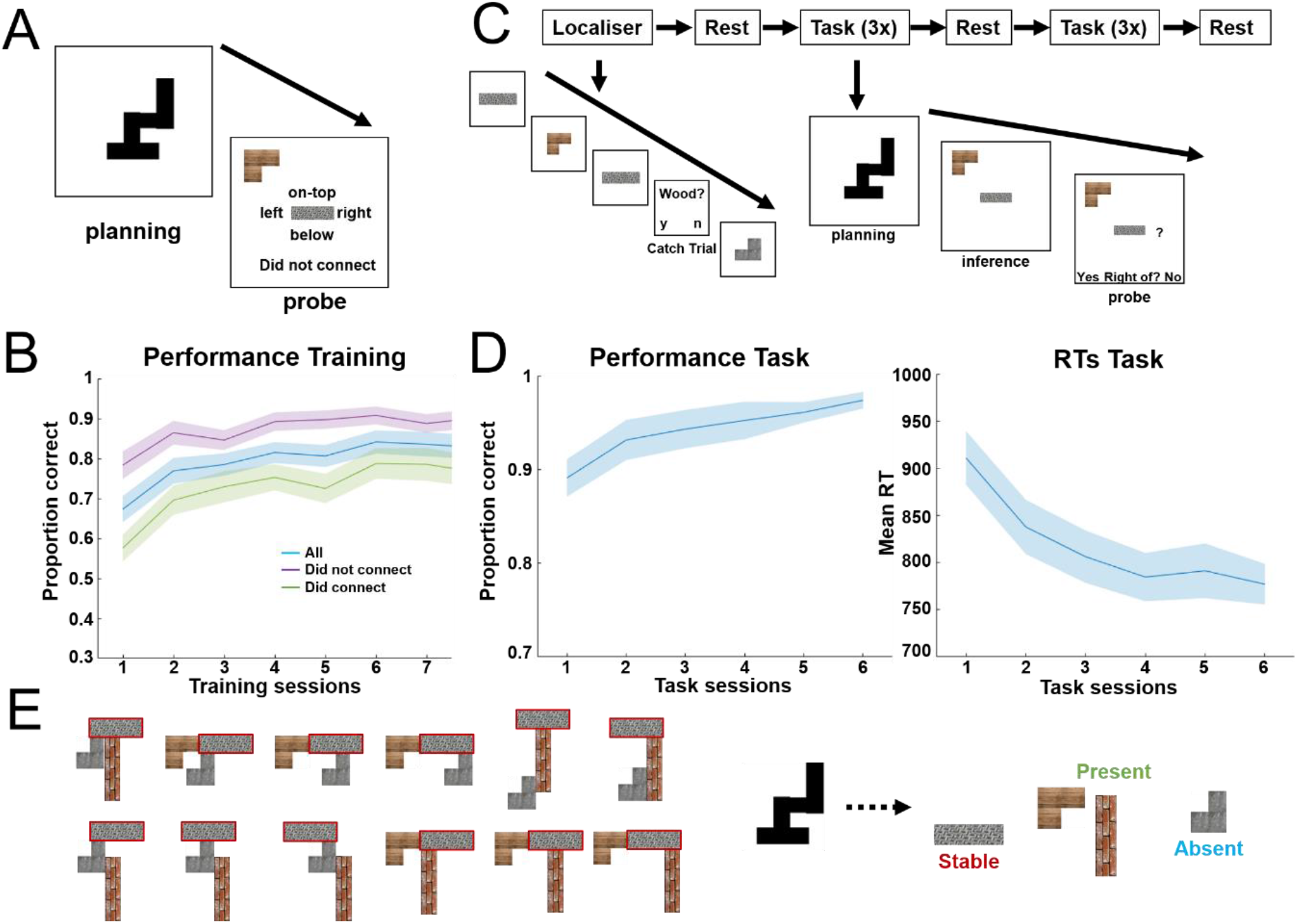
MEG task (A) Subjects were trained on the construction task for two consecutive days. The task consisted of a ‘planning’ and ‘probe’ phase. During planning, subjects were presented with a silhouette and had to make a plan how to construct this silhouette. During probe, subjects were presented with two building blocks and were asked to indicate the relation between these two building blocks in the previous silhouette, if any (‘Was the building block now presented in the top left corner on-top, right, below, or left of the building block now presented in the middle, or did they not connect?’). (B) subjects’ performance on the task improved over time. Performance in trials with unconnected building blocks was generally higher because these trials could often be answered correctly by identifying a probe building block that was not present in the silhouette. (C) After two consecutive days of behavioural training, subjects participated in the MEG experiment. The experiment started with a functional localiser, where subjects observed individual building blocks on the screen and were asked to focus on their texture (wood, concrete, steel or bricks). Intermittently, they received a probe question (‘Was the previous building block made of wood/concrete/steel/bricks?’). The functional localiser was followed by a rest session, followed by three task sessions. The task was identical to the training, except that we included an additional ‘inference’ time window in which subjects were asked to infer the relation between two building blocks, but could not yet indicate a response. In the MEG task, subjects only received probes about building blocks that were connected in the previous silhouette. In the subsequent probe window, subjects were presented with one of four relations (on-top, right, below, or left) and had to indicate whether this was the relation they had inferred. The three task sessions were followed by another rest, followed by another three task sessions and a final rest session. (D) Subjects’ performance again improved over time, such that the proportion of correct responses increased and reaction times decreased with ongoing task experience. (E) In the MEG experiment one building block was always present in every silhouette (*stable*, highlighted in red for an example stimulus set, see Supplementary Figure 4 for all used stimuli), whereas two out of the remaining three had to be inferred (*present*) and one building block was *absent*.

After two days of training in which performance gradually improved (Figure 4B), subjects participated in the MEG experiment (see Figure 4C). The MEG task started with a functional localiser to train binomial classifiers on the individual building blocks (see below). This was followed by six task sessions in total, where subjects played the same task as during training on (initially) novel silhouettes. Every trial in the task had three phases: ‘planning’, ‘inference’, and ‘probe’. During the planning phase, subjects were presented with a silhouette and had to make a plan how to construct this silhouette. During the inference phase, subjects were presented with two building blocks out of the previous silhouette and had to infer the relation between these blocks (‘In your plan, was the building block now in the top left of the screen on-top, right, below, or left of the building block now in the middle of the screen?’). Finally, in the probe phase, subjects were presented with one of four possible relations (on-top, right, below, left) and had to indicate whether that was the relation they inferred. Subjects’ earnings were again dependent on their performance in these probe questions. At the beginning, middle and end of the experiment subjects also underwent a short rest session. During the experiment, subjects displayed high accuracy in their performance (mean reaction time: 836ms, proportion correct: 0.92) with a substantial improvement over time (Figure 4D).

Our first analysis examined the representational content in the MEG sensors during the planning phase. As in the fMRI, we hypothesized that the MEG sensors would contain representations related to the visual appearance of the silhouette, but also to the relational configuration of the inferred building blocks. Because of the always-present block, and because every silhouette has 3 building blocks, we could not perform the perfectly-controlled algebraic analysis (where target and control are comprised of the same blocks in different relational positions). However, a proxy for this analysis is to test whether representational similarity across silhouettes is predicted by how many times the same building block appears in the same relational position in the two stimuli (Figure 4A). We performed RSA over time in the MEG data (see Luyckx et al., 2019 and methods for details). For every trial and at any given time point, we assessed the empirical similarity of sensor representations. We regressed this empirical similarity matrix against predictions from the configural representation and from the visual similarity of the silhouettes (size and shape overlap – Figure 5B). Because we performed this analysis in a multiple regression, the visual regressors also acted as basic controls for the configural regressor. From 200-1000ms post-stimulus onset, there were strong independent effects of all 3 regressors in the MEG signal (particularly for shape and configural representations, Figure 5). Whilst not as cleanly controlled as the fMRI data above, this suggests that the MEG data are also sensitive to both the visual and configural representations.

**Figure 5.**
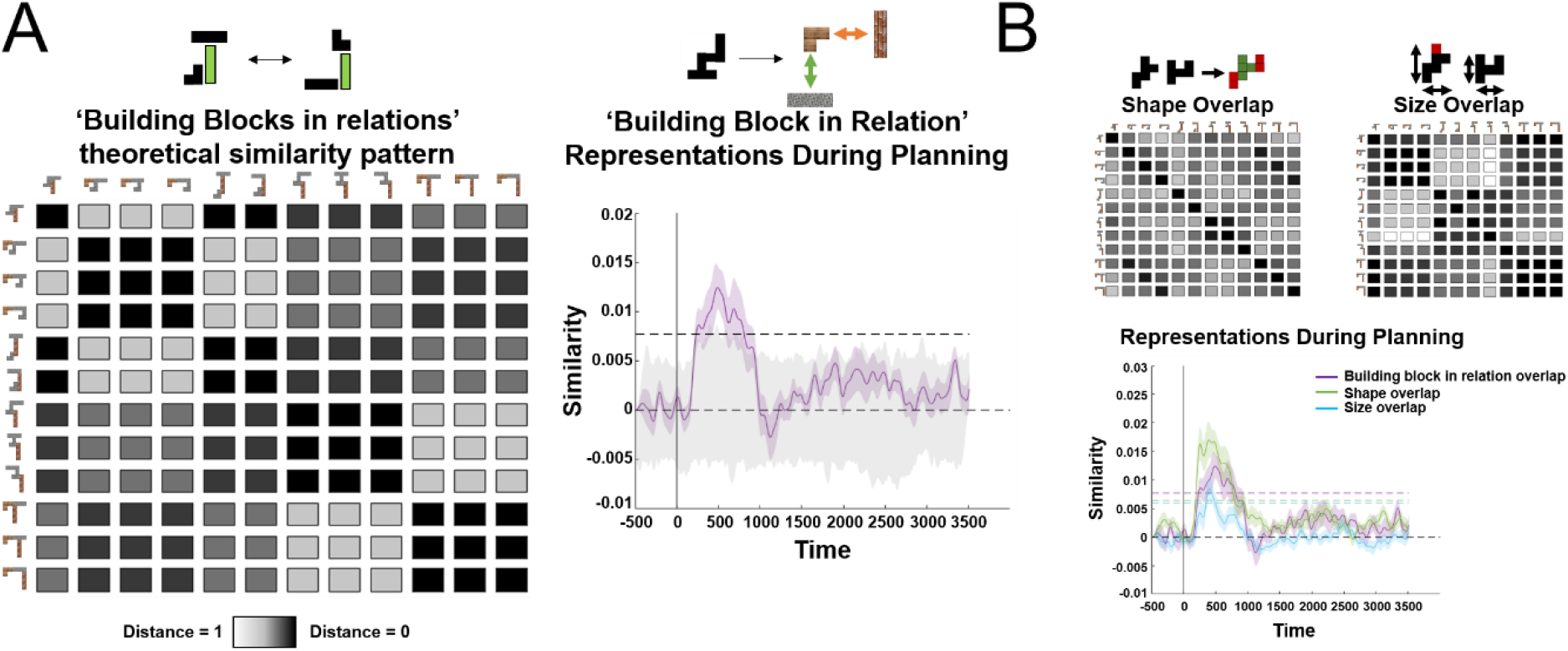
Conjunctive representations akin to the ‘silhouette algebra’ from Figure 3B over time using RSA. (A) Left: We defined a theoretical similarity reflecting the overlap of building blocks in specific relations across silhouettes, and tested whether this similarity predicts empirical similarities of MEG sensor patterns across trials and time-points. Right: We found a significant conjunctive representation, that is representations of silhouettes according to their constituent building blocks in specific relations, during a confined time-window of 200-1000ms in the planning phase (significance assessed using a nonparametric permutation test, see methods for details). (B) We also found effects for shape (pixel) and size representational overlap during a similar time window in planning but with a slightly earlier onset. Note that the purple line in A and B are the same.

### Rapid neural sequences during visual understanding

Next, we asked whether replay may play a role in visual understanding. That is, whether hypothesised visual constructions could be seen in rapid sequences in the MEG data. Recent work has shown that it is possible to measure replay in human MEG data (Kurth-Nelson et al., 2016; Liu et al., 2019). In this approach, classifiers are trained to identify different building blocks and then used to generate predictions for class (re-)activations in independent time-series data, such as during planning periods of the task. This allows to test for reliable transitions between those representations at certain time-intervals, a measure called ‘sequenceness’ thought to be reflective of neural replay. For example, recent studies have shown that when planning a trajectory through a discrete state space (Kurth-Nelson et al., 2016) or resting after learning a sequence of pictures (Liu et al., 2019), individual items are replayed in sequences with a 40ms time-lag, reminiscent of sharp wave ripple activity (Buzsáki, 2015; Diba & Buzsáki, 2007b).

We trained classifiers on building block identity using the functional localiser data in the beginning of the experiment. In line with previous reports, we found that class identifiability peaked at 200ms after stimulus onset (Figure 6A left and middle) and the classifiers displayed high specificity for identifying the correct building block when trained at that time (Figure 6A right-see methods for details on classifier training). We then used these classifiers - trained on the localiser data – to identify reactivations of these representations during planning. We used linear modelling (Liu et al., 2020) to test whether these reactivations occurred in specific (pairwise) orders and at specific time-lags.

**Figure 6.**
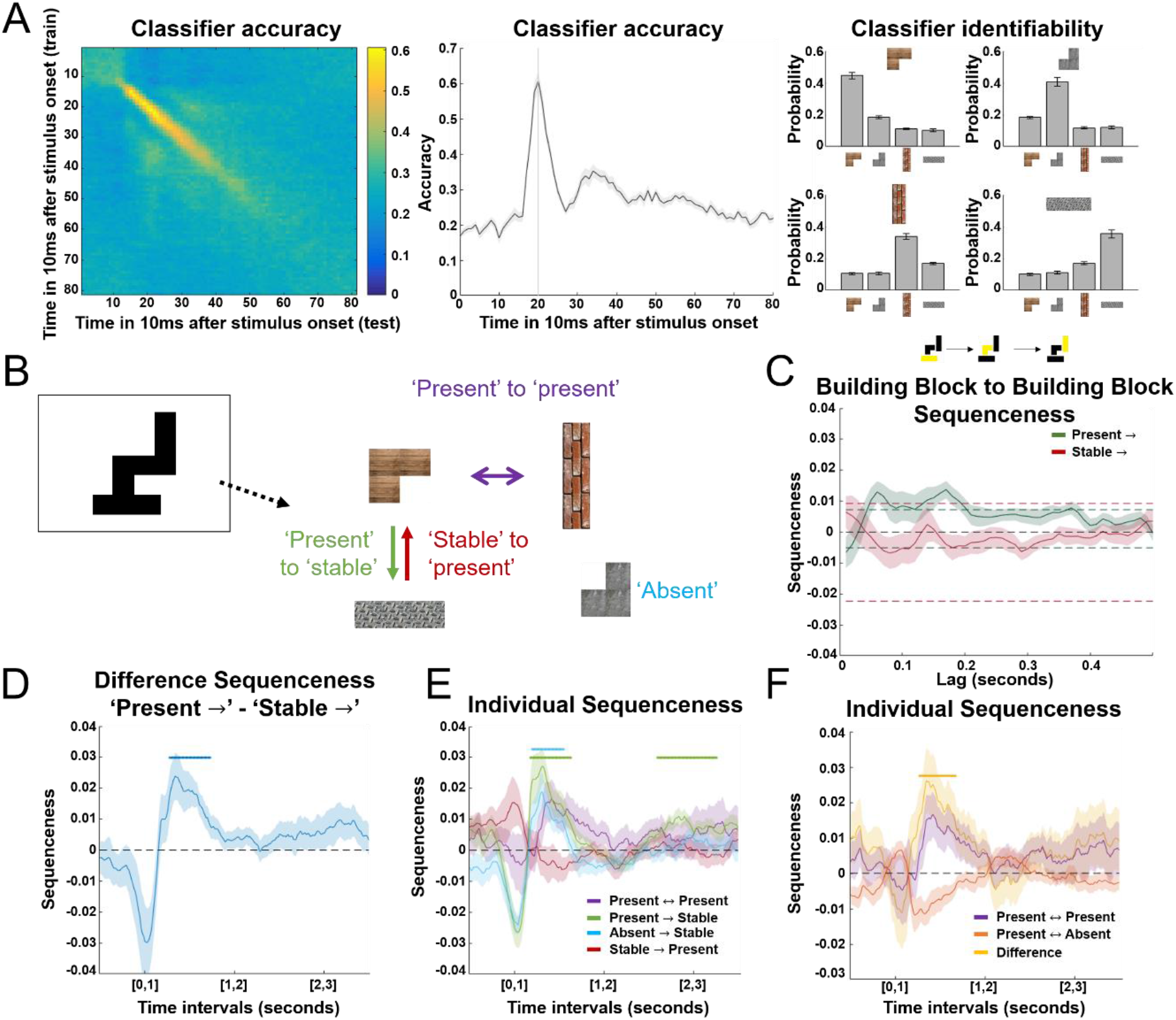
Neural replay underlying visual construction (A) We trained binomial classifiers on the four different building blocks using the data from the functional localiser. We found peak decoding accuracy at 200ms (left and middle) and high class identifiability for the different building blocks for the classifiers trained at 200ms (right). Note the slightly higher confusability of corner and straight building blocks among each other. (B) In every silhouette, one building block was *stable* across silhouettes, two additional building blocks were *present* and one building block was *absent*. This allowed us to define different types of sequences to (green) and from (red) the stable building block as well as between the present (purple) and absent (cyan) building blocks. (C) We investigated effects of neural replay for sequences starting either from the stable or the present building blocks. We found a short (non-significant) predominance of sequences starting from the stable building block for very early lags, followed by a predominance of sequences starting from the present building blocks at later lags with pronounced peaks at 60ms and 170ms. (D) We next investigated the difference between these two types of sequences for different time-intervals of the planning period, and found a brief early predominance of replay starting from the ‘stable’ building block followed by a predominance of replay starting from the ‘present’ building blocks (280-1730ms) during planning. (E) We assessed the individual contributions of the different types of neural replay to these differences and found an unspecific predominance of sequences from both ‘present’ (180-1630ms) and ‘absent’ (200-1550ms) building blocks to the ‘stable’ building block early during planning, and a specific effect from ‘present’ to the ‘stable’ building block late (1600-3260ms) in planning shortly before subjects entered the decision phase of the task. (F) We found a selective predominance of replay between present building blocks over replay between present and absent building blocks in a time window between 270-1670ms.

Importantly, one (***stable***) building block was present in every silhouette. This meant that subjects could use this knowledge to constrain possible configurations. Each silhouette also used two from the remaining 3 building blocks. On each trial, these two (***present)*** building blocks were different, and arranged in different configurations. This left out one (***absent***) building block in every trial that was not present in the silhouette. In some trials the stable block was connected to both present blocks. In other trials the stable block was connected to one of the present blocks and there was also a connection between the two present blocks (Figure 6B - see also Supplementary Figure 4B and C).

To establish whether neural sequences exist, we first examined sequenceness from **stable** and **present** blocks to their connected neighbours in each trial’s silhouette. We initially focussed on the planning period after the first 500ms to be certain to avoid contamination from basic visual processes. We found little evidence for sequenceness starting from the ***stable*** building block, but strong evidence of sequenceness starting from ***present*** building blocks (multiple-comparison-corrected permutation test, Figure 6C – see methods). Note that the x-axis in Figure 6C is the temporal lag – the time difference *between representations that form a sequence*. This effect was significant at a broad range of temporal lags between 30 and 200ms, but had pronounced peaks at 60ms and 170ms – two time-lags that correspond to reports in the previous human replay literature (Kurth-Nelson et al., 2016; Liu et al., 2019, 2021).

To understand the computations executed in replay, we needed to examine how the content of replay changed throughout the planning period. We therefore designed a moving-window analysis. To do this, we averaged over (10ms-200ms) temporal lags shown in Figure 6C. We computed this average sequenceness in 1-second windows centred at every 10ms in the planning period. Hence, unlike Figure 6C, the x-axes in figure 6D-F refer to the time *in the planning period*, not the temporal lag within the sequence. To demonstrate this method, Figure 6D shows the difference between sequences that start with ***present*** compared to those that start with ***stable***. This is the same difference shown in Figure 6C but now measured at different times in the planning period. Here, the effect in Figure 6C is revealed as a significant cluster covering the time-range 280-1730ms (cluster corrected p< 0.05 - see methods).

Also of note is a strong early negative effect (see also Supplementary Figure 5). This might indicate a short very early period where replay emanates from the stable blocks, but more likely is a confound: Unlike all other blocks, the stable block can be represented **before** the onset of the stimulus (as it is there in every trial). Any later activation of other blocks (when the stimulus appears) will be measured as a forward sequence (as it comes after the pre-stimulus representation **stable**). We therefore refrain from interpreting this peak here and in all later graphs.

### Replay as generative hypothesis construction

Our task structure confers a clear optimal strategy for sequential hypotheses, as the stable block constrains what solutions are possible. Subjects should begin the construction process by testing the **stable** block with all other candidate blocks. Once the neighbour(s) of the stable block is (are) resolved, the final step is to resolve any remaining connections between the two **present** blocks. Replay followed exactly this progression (Figure 6E, 6F). The earliest sequences in the planning period all proceeded towards the **stable** block and did not distinguish between **present** (180-1630ms) and **absent** (200-1550ms) blocks (blue and green lines in figure 6E – note that these two analyses are orthogonal, the blue and green lines are independent measurements). Sequences that did not involve the **stable** blocks emerged later (270-1670ms) and *only involved the* ***present*** *blocks* (Figure 6F – yellow line. Note **present**<->**present** sequences in purple are shown in 6E and 6F to allow visual comparison of timings.). Finally, at the end of the planning period sequences to the stable block re-emerged, but only those involving the **present** blocks (1600-3260ms, Figure 6E).

Taken together, these results indicate a role for replay in constrained hypothesis generation during visual understanding. Replay followed the optimal strategy for hypothesis generation, starting with unspecific sequences to the stable block, proceeding to infer connections between present blocks, and converging on sequences that only include correct blocks.

## Discussion

The hippocampal formation and prefrontal cortex are critically involved in visual understanding (Aly et al., 2013; Córdova et al., 2019; Ruiz et al., 2020), the instantiation of a cognitive map during spatial and conceptual navigation (Constantinescu et al., 2016; Garvert et al., 2017; Hartley et al., 2014; O’keefe & Nadel, 1978; Tolman, 1948), and model-based planning in RL (Brunec & Momennejad, 2021; Miller et al., 2017; Stachenfeld et al., 2017; Vikbladh et al., 2019). A key problem underlying these functions is learning an efficient representation of the state space that deals with the considerable complexity of naturalistic problems and enables the generalisation of knowledge to novel instances. Motivated by insights into the neural organisation underlying efficient spatial navigation, we developed a novel visual construction task that allowed us to probe whether compositional neural representations subserve flexible visual understanding. In two independent studies, we found evidence for such compositional representations that allow to flexibly combine knowledge about basic building blocks and abstract task structure to form novel conjunctive representations. We further identified neural replay, thought to underlie learning and inference in the HC-PFC circuit, as a candidate mechanism of generative hypothesis testing.

We probed the neural representations that facilitate the generalisation of knowledge in previously unseen visual construction problems. We found evidence for a neural code that reflects the generalisable embedding of individual building blocks into a certain relational position across different silhouettes, such as a building block ‘on-top’ of another building block. Using MEG, we found these representations during a confined time-window of 200-1000ms after stimulus onset underlying visual construction. Using fMRI, we found that these ‘conjunctive’ codes accurately predicted representations in mPFC and anterior hippocampus. The rich stimulus set of the fMRI task allowed us to define a ‘silhouette algebra’, which implies the encoding of individual building blocks embedded in specific relational positions that generalise across different silhouettes. This ‘silhouette algebra’ is closely aligned to a ‘scene algebra’ that was recently proposed in a generative model of scene understanding (Eslami et al., 2018). This model proposed a compositional basis of visual understanding, where certain object-features (like ‘blueness’ or ‘roundness’) in a given scene could be predicted by representations of those object-features in other scenes. Our results provide evidence for a biological implementation of a compositional account of scene understanding in the hippocampal-prefrontal circuit. Further, these ‘silhouette algebra’ effects in anterior hippocampus are highly overlapping with recent evidence that activity patterns in this area can be used to predict representations of stimuli by other stimuli and their translation in an abstract stimulus space (Morton et al., 2020).

These results are closely related to prior work highlighting the importance of mPFC and hippocampal representations in the construction of novel compounds, such as ‘tea jelly’, out of known compounds, such as ‘tea’ and ‘jelly’ (Barron et al., 2013; Bongioanni et al., 2021). We indeed detected representations in mPFC that reflect the use of a building block in a given compound irrespective of its relational embedding, highly overlapping with ‘tea jelly’ representations reported earlier (Barron et al., 2013). Such representations are predicted under a ‘factorised code’ (Behrens et al., 2018; Manns & Eichenbaum, 2006), i.e., a representation of basic sensory building blocks that can be flexibly combined with structural knowledge to form novel conjunctive representations. It is an important challenge to understand how such structural or relational knowledge itself is represented efficiently, such that it can be flexibly inferred (Kemp & Tenenbaum, 2008) and adjusted to novel contexts, akin to a basis set for structural reasoning (Mark et al., 2020).

Based on the involvement of the hippocampal-prefrontal circuit in the present visual construction task, we asked whether generative replay provides a candidate mechanism underlying flexible construction. It has been hypothesised that replay reflects sampling from a generative model of the world to facilitate inference (Foster, 2017), enable generalisation (Barry & Love, 2021), and train a recognition model (Dayan et al., 1995), thus providing a key mechanism for active hypothesis testing about possible futures an animal might encounter. Here, we tested whether replay is also involved in hypothesis testing about possible presents, i.e., correct ways of de-constructing a silhouette.

In a MEG study, we detected such generative replay underlying constructive hypothesis testing. We introduced biased silhouettes in which one building block was *stable* (always present) and two building blocks had to be inferred (*present*). Consequently, we distinguished neural sequences that start from the ‘stable’ or ‘present’ building blocks during planning. We found a predominance of sequences starting from the ‘stable’ at the very beginning of planning, followed by a predominance of sequences starting from the ‘present’ building blocks, with the latter type of sequenceness peaking at a time-lag of 60ms and 170ms. A closer examination of the time course of these sequences revealed an unspecific predominance for sequences towards the stable building block from both the present and absent building block early during planning, but a specific effect for replay from the correctly inferred present building blocks towards the stable building block late during the planning period just before subjects enter a decision phase. Importantly, we also detected an emerging predominance for sequences *between* correctly inferred present building blocks over sequences between present and absent building blocks as planning proceeds.

This provides important insight into the neural mechanisms that subserve efficient planning in the present paradigm that is both sensitive to experienced task structure (‘start with the stable building block’) and the specifics of a visual target (‘now find the other present building blocks in contrast to the absent building block’). The present results suggest that generative replay might underlie this process of hypothesis testing, which becomes increasingly refined as planning proceeds. These results are in line with previous reports suggesting a role for replay when planning (Kurth-Nelson et al., 2016) and learning (Schuck & Niv, 2019) trajectories through an abstract state space in a RL problem and evidence from recordings in animals suggesting that replay can explore novel trajectories (Gupta et al., 2010; Ólafsdóttir et al., 2015). Our findings are in line with notions that generative replay provides a mechanism for efficiently learning and sampling from a generative model of the world (Foster, 2017), in line with a crucial role of replay in planning and RL (Antonov et al., 2021; Mattar & Daw, 2018) but also structure learning (Evans & Burgess, 2020). In the present study, we provide evidence for replay as a mechanism for testing hypotheses underlying compositional understanding.

In conclusion, we developed a paradigm to probe the neural mechanisms that underlie efficient state space representations in sophisticated visual understanding problems. In close alignment with neural representations subserving both navigation and model-based RL, we found conjunctive representations in the hippocampal formation and prefrontal cortex that flexibly generalise knowledge about relations between objects in a scene. Further, we identified generative neural replay as a candidate mechanism underlying gradual hypothesis testing in visual construction problems. Together, these results provide insight into efficient neural representations that enable flexible generalisation, raising the intriguing hypothesis of a shared neural code underlying navigation, model-based RL, and visual understanding based on a cognitive map of task structure.

## Methods

### fMRI task

#### Participants

30 subjects (25 females, mean age: 22.9, range: 19-33) participated in the behavioural training and subsequent fMRI experiment. Additionally, we scanned two pilot subjects and one subject did not participate in the fMRI part of the experiment after the behavioural training. All subjects were recruited from the UCL psychology subject pool, had no history of neurological or psychiatric illness and had normal or corrected-to normal vision. All subjects gave written informed consent and the study was approved by the UCL ethics committee (ethics code: 11235/001).

#### Training and fMRI task

Subjects completed two tasks during behavioural training on two consecutive days. Initially, subjects completed four sessions (50 trials each) of the construction task on the first day of training. In this version of the training, there was no time restriction, and subjects were supposed to familiarise themselves with the task contingencies. In every trial, subjects were presented with the nine basic building blocks at the top of the screen and saw a target silhouette at the bottom left. They then had to construct the target silhouette by selecting the correct building blocks and moving them around on the screen using a computer keyboard, being instructed that a construction would only be marked correct if they find a solution with the minimum number of elements. Further, every building block could only ever be used once. Silhouettes increased in size and complexity over the course of this training. On the second day of training, subjects had to solve five sessions (70 trials each) of a similar task but this time only select the correct building blocks without actually constructing the silhouette. This version of the training had a time restriction, such that subjects had 6 seconds to infer a construction plan for a given silhouette followed by 6 seconds to select the correct building blocks. This task was designed to train subjects on the rapid mental construction of a silhouette that was required in the fMRI. In both tasks, subjects received feedback at the end of a trial indicating whether the construction or selection was correct, and they received 3 pence per correct answer in the second version of the training task.

To impose hierarchical learning, we gradually introduced hierarchical building blocks into the training regime. In the first two training sessions of training day 1, subjects only had to construct silhouettes consisting of two building blocks. 27 of these 50 silhouettes in each session were hierarchical building blocks (3 trials per hierarchical building block) as illustrated in Figure 2B (second row). In the next two sessions of the construction task, subjects received larger silhouettes that often contained one or two hierarchical building blocks (third session: 18 hierarchical building blocks, 18 silhouettes with one hierarchical building block and one extra basic building block; fourth session: 18 silhouettes consisting of two hierarchical building blocks and 18 silhouettes with one hierarchical building block and one extra basic building block). Of the 70 trials in every session of the second training task, 24 where silhouettes that consisted of two hierarchical building blocks.

In the fMRI experiment, subjects had to solve a similar task to the second training task. Here, subjects were presented with a silhouette for 2 seconds and were tasked to mentally construct this silhouette. In 90% of trials, this was followed by a fixation cross for 1 second before presenting the next silhouette. In 10% of the trials, the silhouette was followed by a probe trial. In this probe trial, subjects were shown one or two basic building blocks and asked whether this/these building block/s can be used for the construction of the previous silhouette. Subjects had 2 seconds to respond ‘yes’ or ‘no’ via button press and received 20 pence for every correct answer. Every session in the scanner consisted of 288 trials in total, and subjects completed three sessions. In half of these trials, subjects were probed on a silhouette that either consisted of two basic or hierarchical building blocks (two repetitions per silhouette), combined with ‘on-topness’ or ‘besideness’ (i.e., one building block is on-top or left/right of another building block). In the other half of the trials, subjects were presented with one of the nine basic or hierarchical building blocks (eight repetitions per building block). In order to minimise effects of visual overlap of individual building blocks with silhouettes using these building blocks on the screen, the individual building blocks were presented at various locations throughout a session (twice at the top/bottom/left/right of the screen).

After the fMRI task, we assessed subjects’ individual similarity judgements about silhouettes that were presented in the scanner. To do so, subjects completed two sessions consisting of 120 trials in total, where they were presented with a target silhouette in the top middle of the screen and had to judge whether this target silhouette was more similar to a silhouette presented at the bottom left or right. Subjects had 6 seconds to make this judgement, followed by a 1 second inter-stimulus interval. In half of these 120 trials subjects were probed about silhouettes using basic building blocks and half of trials consisted of silhouettes using hierarchical building blocks. In the first of these two sessions subjects were instructed to focus on visual similarity, while in the second session subjects were instructed to focus on ‘construction similarity’ (‘which silhouette is more similar in terms of how you would construct them?’).

#### fMRI data acquisition

FMRI data was acquired on a 3T Siemens Prisma scanner using 32 channel head coil. Functional scans were collected using a T2*-weighted echo-planar imaging (EPI) sequence with a multi-band acceleration factor of 4 (TR = 1.450 s, TE = 35 ms, flip angle = 70 degrees, voxel resolution of 2×2×2mm). A field map with dual echo-time images (TE1 = 10ms, TE2 = 12.46ms, whole-brain coverage, voxel size 2×2×2mm) was acquired to correct for geometric distortions due to susceptibility-induced field inhomogeneities. Structural scans were acquired using a T1-weighted MPRAGE sequence with 1×1×1mm voxel resolution. We discarded the first six volumes to allow for scanner equilibration.

#### Pre-processing

All pre-processing steps and subsequent imaging analyses were performed with SPM12 (Wellcome Trust Centre for Neuroimaging, http://www.fil.ion.ucl.ac.uk/spm). Functional images were corrected for signal bias and realigned to the first volume in the sequence using a six-parameter rigid body transformation to correct for motion. Images were then spatially normalised by warping subject-specific images to MNI (Montreal Neurological Institute) reference coordinates and smoothed using a 6-mm full-width at half maximum Gaussian kernel. The RSA-analysis was performed on unsmoothed data before smoothing the resulting contrast estimates (see below).

#### Repetition suppression analysis

We employed univariate repetition suppression analysis to test for compositional representations of individual building blocks within a silhouette. To do so, we modelled the onset of all objects on the screen as stick functions, and defined several parametrically modulated regressors of interest and to control for potential confound variables. In total, we defined four control regressors that account for repetition suppression due to size or pixel non-overlap and change in the number of building blocks in a silhouette. Size non-overlap was defined the absolute difference in height and width of visual silhouettes, and pixel non-overlap as the maximum proportion of overlap of pixels of two silhouettes relative to their full ‘pixel-size’ subtracted from 1. We also added the number of building blocks in a silhouette as additional fourth control regressor. The effects for these control regressors are shown in Figure 2. Next, we defined three building block non-overlap regressors that account for compositional representations, i.e., representations of individual building blocks within a silhouette. We defined a regressor that reflected the proportion of non-overlap of the basic building blocks in a present silhouette with the basic building blocks of the previous silhouette (see supplementary information for an illustration). This regressor only had a unique value for small silhouettes that did not consist of hierarchical building blocks. This is because there was more than one solution of basic building blocks in large silhouettes (built with two hierarchical building blocks and four basic building blocks). Consequently, we split up this regressor that reflected the non-overlap of basic building blocks into trials with small (two basic building blocks) and large (two hierarchical building blocks, four basic building blocks) silhouettes. For large silhouette-trials that had more than one basic building block solution, we computed the average of building block non-overlap weighted by the different solutions for a given silhouette. In addition to those basic building block non-overlap regressors, we defined a hierarchical building block non-overlap regressor following the same logic but with hierarchical building blocks. Just as the regressor for basic building block non-overlap in small silhouettes, the regressor for hierarchical building block non-overlap (in large silhouettes) had only unique solutions. Finally, we defined three regressors that accounted for relational non-overlap. These regressors differentiated between trials of silhouettes that used the same or a different relational operation (putting a building block on-top or beside another building block) compared to the previous trial. We split this regressor into trials of small silhouette transitions (relational operation for basic building blocks), large silhouette transitions (relational operation for hierarchical building blocks), and transitions between small and large silhouettes. In order to make all these parametric regressors comparable, they were projected onto an interval ranging from -1 to 1.

Because of the sensitivity of the blood oxygen level-dependent (BOLD) signal to motion and physiological noise, all GLMs also included six motion regressors and their derivatives obtained during realignment, as well as 6 regressors for cardiac phase, 6 for respiratory phase and 2 for respiratory volume extracted with an in-house developed Matlab toolbox (Hutton et al., 2011). Sessions were modelled separately within the GLMs.

To obtain the ‘tea jelly’-like compositional representation results (Figure 3C), we combined the effects for building block non-overlap with unique solutions, i.e., the basic building block non-overlap in small silhouettes and the hierarchical building block non-overlap.

#### ‘Silhouette algebra’ analysis

To test for the presence of a conjunctive code, we assessed the representational distance between algebra terms, target silhouettes and reference silhouettes. We performed volumetric searchlight RSA (Kriegeskorte et al., 2008) based on a GLM approach similar to (Hunt et al., 2018). Effectively, we asked whether we can predict empirical distances between activity patterns for algebra terms, target silhouettes, and reference silhouettes by theoretical distances predicted by a conjunctive code whilst controlling for visual confounds based on size and shape overlap. To do so, we first obtained individual coefficient estimates for all stimuli used in the fMRI task based on a first-level univariate GLM on unsmoothed data. We then defined searchlights across every voxel including the 100 cortical voxels with smallest geodesic distance from the central voxel (Kriegeskorte et al., 2006). Coefficient estimates in every searchlight were pre-whitened. Both the searchlight definition and pre-whitening were based on adapted scripts from the RSA toolbox (Nili et al., 2014). We then defined a representational distance metric (defined as 1-correlation) between algebra terms, target and reference silhouettes. To do so, we first computed all possible algebra terms as shown in Supplementary Figure 3. We then computed the distance to the respective target and reference silhouette, resulting in a sparse representational matrix of size [(algebra terms + target silhouettes + reference silhouettes) x 3 sessions] x [(algebra terms + target silhouettes + reference silhouettes) x 3 sessions]. To avoid any within-session similarity effects, we only computed and compared cross-session distances. Next, we tested whether this empirical representational distance matrix could be predicted by a theoretical representational distance that reflects a conjunctive code as shown in Figure 3B. For every searchlight, we computed a GLM to assess the prediction of the empirical representational distance matrix based on a conjunctive representation, whilst controlling for two additional theoretical distances based on the shape and size of the objects. This ensured that any shared variance between conjunctive and visual confound representations was removed, and resulted in a single conjunctive representation map per subject. These coefficient maps were smoothed using a 5mm FWHM kernel in line with a recent study based on a similar approach (Baram et al., 2020).

#### Multiple comparison correction

To assess statistical significance on the group-level for conjunctive representations, we performed family-wise error (FWE) corrected sign-flip permutation tests (Nichols & Holmes, 2002) using PALM (Winkler et al., 2014) either using a pre-defined ROI of the hippocampal formation based on the Juelich anatomical atlas (Eickhoff et al., 2005, 2007) or in a more exploratory whole-brain approach. Coefficient values of every subject were randomly multiplied by 1 or -1 based on the null-hypothesis that these coefficient values are symmetrically distributed around 0. To create a null distribution of the means this process was repeated 5000 times, and the true value was then compared to this null distribution. On the whole brain level, we used a maximum cluster mass statistic (Nichols & Holmes, 2002) for FWE correction based on a cluster forming threshold of p < 0.001.

### MEG task

#### Participants

20 subjects (15 females, mean age: 25.4, range: 20-36) participated in the behavioural training and subsequent MEG experiment. We scanned two pilot subjects prior to the experiment and one subject had to be excluded from the analysis due to impaired vision. All subjects were recruited from the UCL psychology subject pool, had no history of neurological or psychiatric illness and had normal or corrected-to normal vision. All subjects gave written informed consent and the study was approved by the UCL ethics committee (ethics code: 11235/001).

#### Training and task

Subjects completed two tasks during behavioural training. Initially, subjects completed two sessions of a construction task (50 trials each) of the same structure as in the beginning of the training for the fMRI task. In this task subjects only had four different building blocks available to construct silhouettes. After two sessions of the construction task, subjects were trained on a second version of the task that required them to make judgements about the relational configuration of given silhouettes. Subjects saw a silhouette for 6 seconds and had to infer the relational positions of individual building blocks in the silhouette. This was followed by a question screen lasting for 6 seconds, in which subjects were shown two building blocks and asked how they related to each other in the previous silhouette. Specifically, one of these building blocks was presented in the middle of the screen and the other at the top left of the screen, and subjects had to infer whether the building block in the top left was on-top, right, below, or left of the building block in the middle of the screen. They also had the option to indicate that the two building blocks did not connect in the previous silhouette. Subjects completed 3 sessions of this task on the first day and 5 sessions on the second day of training at received 5 pence for every correct answer.

After being trained on the task for two consecutive days, subjects participated in an MEG experiment on the day after the second day of training. In the scanner, subjects starting with a resting session, in which subjects saw a fixation cross for 4 minutes and were instructed to maintain a state of wakeful rest. This was followed by a localiser screen for individual building blocks, which allowed us to train classifiers to decode individual building blocks from sensor activity (see below). Subjects completed two sessions in which each of the four building blocks was shown 25 times on the screen for 2 seconds. Subjects were instructed to focus on the building block identity, and particularly its texture (bricks, concrete, steel, or wood). To ensure that subjects actively engaged with the task, 10% of trials were followed by probe questions in which subjects had to indicate within 2 seconds via button press whether the previous building block was made of bricks/concrete/steel/wood. These two localiser sessions were followed by three task sessions (48 trials each). Subjects had to perform a task that was very similar to the training task where they had to infer the relation between two building blocks in a previous silhouette. In contrast to the training, the presented building blocks always connected to each other in the previous silhouette, such that the ‘did not connect’ option was removed in the MEG task. In this task, subjects saw a silhouette and had to infer a plan of its construction for 3.5 seconds, followed by a screen showing two building blocks out of the previous silhouette for 3.5 seconds, in which subjects had to infer how one building block related to the other in the previous silhouette. Finally, subjects saw a question screen for 1.5 seconds in which they were presented with one of four possible relations (on-top of, right of, below of, or left of) and had to indicate whether this was the relation they had inferred via button press (‘yes’ or ‘no’). In these question screens, probe relations could either be presented as text written at the bottom of the screen or via a question mark at the corresponding location (on-top, right, below, or left) of the building block presented in the middle of the screen to ensure that subjects process both the semantic meaning and the actual use in the construction of the inferred relation. The three task sessions were followed by another 4-minute rest period, followed by another 3 task sessions and a final rest period.

#### MEG data acquisition and preprocessing

MEG was recorded continuously at 1200 samples/second using a whole-head 275-channel axial gradiometer system (CTF Omega, VSM MedTech), while participants sat upright in the scanner. Subjects indicated ‘yes’ and ‘no’ responses in both the functional localiser and MEG task using a scanner-compatible button box.

The preprocessing protocol closely followed a recently published study (Liu et al., 2019). Data were resampled from 1200 to 100 Hz to improve signal to noise ratio and high-pass filtered at 0.5 Hz using a first-order IIR filter to remove slow drift. Subsequently, an ICA (FastICA, http://research.ics.aalto.fi/ica/fastica/) was performed to decompose the data into 150 temporally independent components and their corresponding sensor topographies. Artifact components were identified using automated inspection based on spatial topography, time course, kurtosis of the time course and frequency spectrum. Eye-blink artifacts can be identified based on high kurtosis (>20) and mains interference based on a low kurtosis and a frequency spectrum dominated by 50 Hz line noise. Based on these definitions, artifacts were rejected by subtracting them out of the data. Epoched data from the functional localiser and planning period during the MEG task was baseline-corrected by subtracting the mean sensor activity 100ms before stimulus onset from the data. Subsequent analyses were performed directly on the filtered, cleaned MEG signal in units of femtotesla on the whole-brain sensor level.

#### RSA

We performed GLM-based representational similarity analysis akin to our fMRI analysis. During the planning and inference period of the task we obtained empirical representational similarity matrices for the different stimuli based on a similar approach reported in (Luyckx et al., 2019). We defined a design matrix that specified the present silhouette at a given trial using one-hot vectors (12 one-hot vectors for 12 silhouettes in total) and one additional regressor to account for the mean activity. Using this design matrix, we then obtained sensor coefficients for each stimulus at any given time point, resulting in a sensor x stimulus x time-point matrix. The coefficients were pre-whitened using an adapted script from the RSA toolbox (Nili et al., 2014) and then used to compute Pearson correlation coefficients between sensors for every individual silhouette for every time-point. This resulted in a representational similarity matrix for silhouettes across time points for both the planning and inference period. Akin to the fMRI conjunctive code analysis, we then specified a GLM predicting these empirical similarities using different (z-scored) theoretical similarities across time (see Figure 5). During the planning period when subjects saw a silhouette on the screen, we defined conjunctive representations (building block in a specific relational position) as well as size and pixel overlap as theoretical representational similarities.

We defined a non-parametric permutation threshold to test for statistical significance. We repeated the above analyses with randomly shuffled predictor representational similarities 5000 times to create a null distribution of predictive coefficients. To define a significance threshold, we defined the 97.5^th^ and 2.5^th^ percentile of the range of the shuffled predictive coefficients as upper and lower significance thresholds.

#### Multivariate Decoding

Training of the classifiers and subsequent sequenceness analyses closely followed previously published approaches (Liu et al., 2019), which are discussed in detail elsewhere (Liu et al., 2020).

We trained 4 Lasso-regularised regression models on the four different building block classes in the functional localiser data, using only sensors that were not rejected in all individual scanning sessions. This provided a decoding model based on a binomial classifier per building block, where we defined all trials in which a corresponding building block was present as positive examples and all other trials as negative examples. To decorrelate the classifiers we also included null data, defined as the sensor data from 500ms before stimulus onset. In line with previous work (Kurth-Nelson et al., 2016; Liu et al., 2019) we found maximum decodability (defined as the highest probability output among all classifiers being assigned to the correct class) of individual building blocks 200ms after their onset and consequently trained the binomial classifiers on that time for the subsequent sequenceness analysis. In line with previous work (Kurth-Nelson et al., 2016), we used and L1 penalty of 0.006, encouraging sparsity in the sensor representations for the individual building blocks.

#### Sequenceness Measure

Please see (Liu et al., 2019, 2020) for a detailed methodological explanation of the sequenceness measure used in the present study. In brief, we used the trained classifiers to obtain class reactivation predictions for the independent planning data. The sequenceness measure is then obtained by applying a GLM approach at two levels. At the first level, we obtain empirical pairwise transitions or sequences between class reactivations by using a linear model to test whether certain stimulus reactivation patterns are predictive of other reactivation patterns at different time-lags (with a maximum lag of 500ms). This results in a 4×4 matrix of empirical building block transitions for every time-lag. Here, we did not include additional nuisance regressors to control for confounding effects, such as an alpha oscillation (Liu et al., 2019), since we applied this analysis to an active planning period rather than a rest phase. At the second level, we then ask whether this pattern of empirical transitions at different time-lags is predicted by theoretical transition matrices whilst controlling for the mean and self-transitions.

We defined three different types of sequences of interest corresponding to three theoretical transition matrices: sequences from the ‘stable’ to the ‘present’ building blocks, sequences from the ‘present’ to the ‘stable’ building blocks and sequences between the ‘present’ building blocks (if applicable – in trials where the ‘stable’ building block was in the middle of the silhouette the two ‘present’ building blocks did not connect). We averaged the latter two to obtain the global effect of sequences starting from the ‘present’ building block as shown in Figure 6C.

Figure 6C shows the effects of the sequenceness analysis when applied to the full planning period except for the first 500ms to allow for basic visual processing. The obtained sequenceness effects were tested against control sequences, where one building block in the true theoretical transition matrix was replaced by the absent building block. This results in two alternative sequences for each of the three sequence types (‘present’ to ‘present’ if applicable) per trial (288 trials in total). We then treated the minimum and maximum of these control sequences across time-points (averaged over trials) as statistical bound, against which we compared the sequenceness for the true sequences.

Figure 6D shows the individual sequences starting from the ‘stable’, ‘present’, or ‘absent’ building blocks for different time intervals using a sliding-window of width 1000ms with a step-size of 10ms, averaged for time-lags 10ms-200ms. Cluster-based statistics was obtained similarly to (Eldar et al., 2018). In the sequenceness time-series, we assessed the length of consecutive time-points exceeding the critical t-value of +/-2.09 (two-sided P value of 0.05 for df=19). The data were then shuffled 10000 times by randomly multiplying half of the subjects’ time-series by -1 and obtaining the maximum length of consecutive time-points exceeding the critical t-value for that shuffle. We then defined the 95^th^ percentile of cluster lengths from the shuffled data as cluster-based significance threshold, against which we tested the original data.

## Acknowledgements

We would like to thank Avital Hahamy for very helpful comments on the fMRI analyses and an earlier version of the manuscript, and Aaron Bornstein, Nicolas Schuck and Peter Dayan for valuable comments on the analyses. We also thank Helen Barron and Mona Garvert for very helpful comments at an earlier stage of the analyses. We thank Nadege Corbin and Martina Callaghan for crucial help with the fMRI data acquisition and preprocessing.

## Author contributions

P.S., S.M., Z.K.N. and T.B. designed the experiments, P.S. collected the data, P.S., A.B., Y.L., T.M., Z.K.N. and T.B. analysed the data, P.S., A.B., L.Y., T.M., S.M., R.D., M.B., Z.K.N. and T.B. interpreted the data, P.S. and T.B. wrote the paper.

**Correspondence and requests for materials** should be addressed to P.S.

## Supplementary information

**Supplementary Figure 1.**
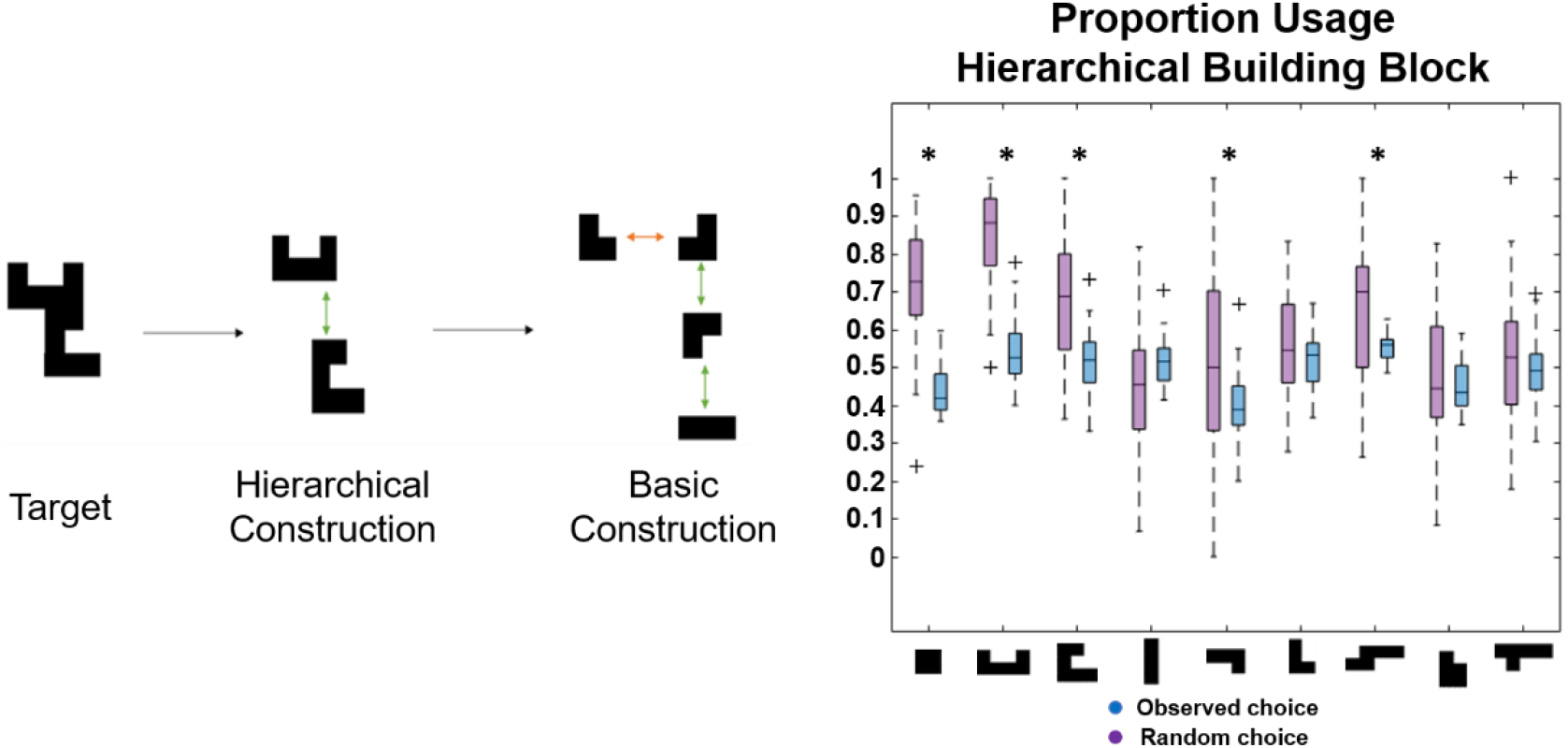
Preferences for ‘hierarchical chunking’ for the individual hierarchical building blocks.

**Supplementary Figure 2.**
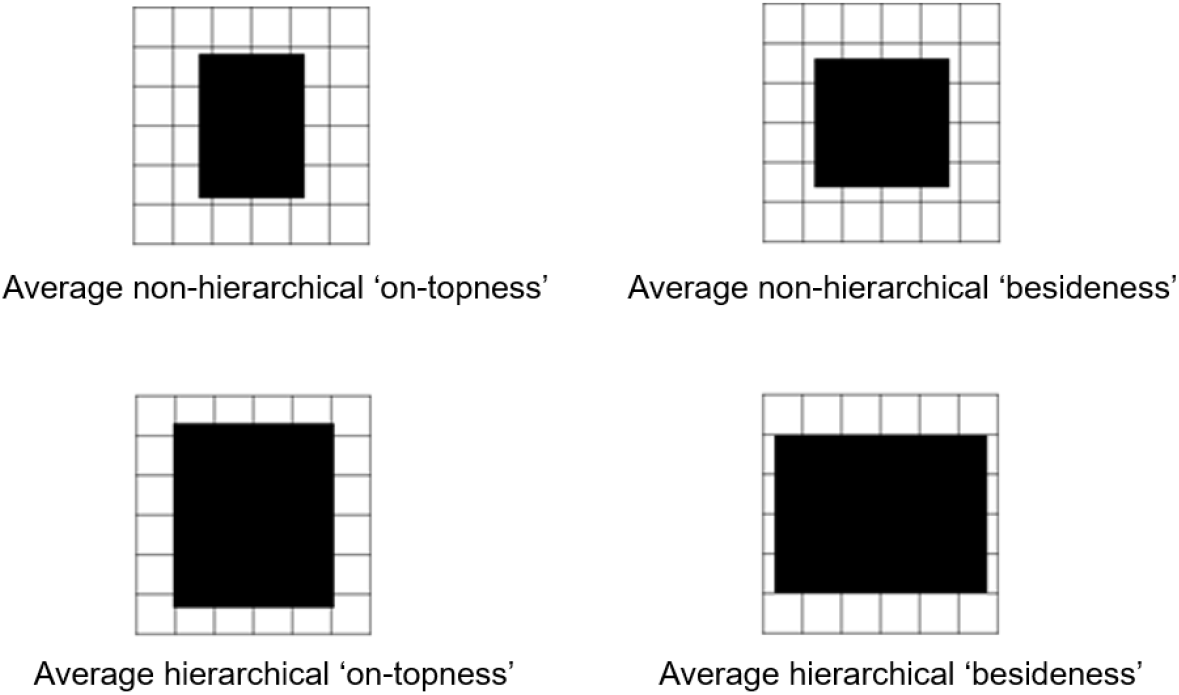
Average size for non-hierarchical (top row) and hierarchical (bottom row) compounds built by placing one (basic or hierarchical) building block on-top (left column) or beside (right column) another (basic or hierarchical) building block.

**Supplementary Figure 3.**
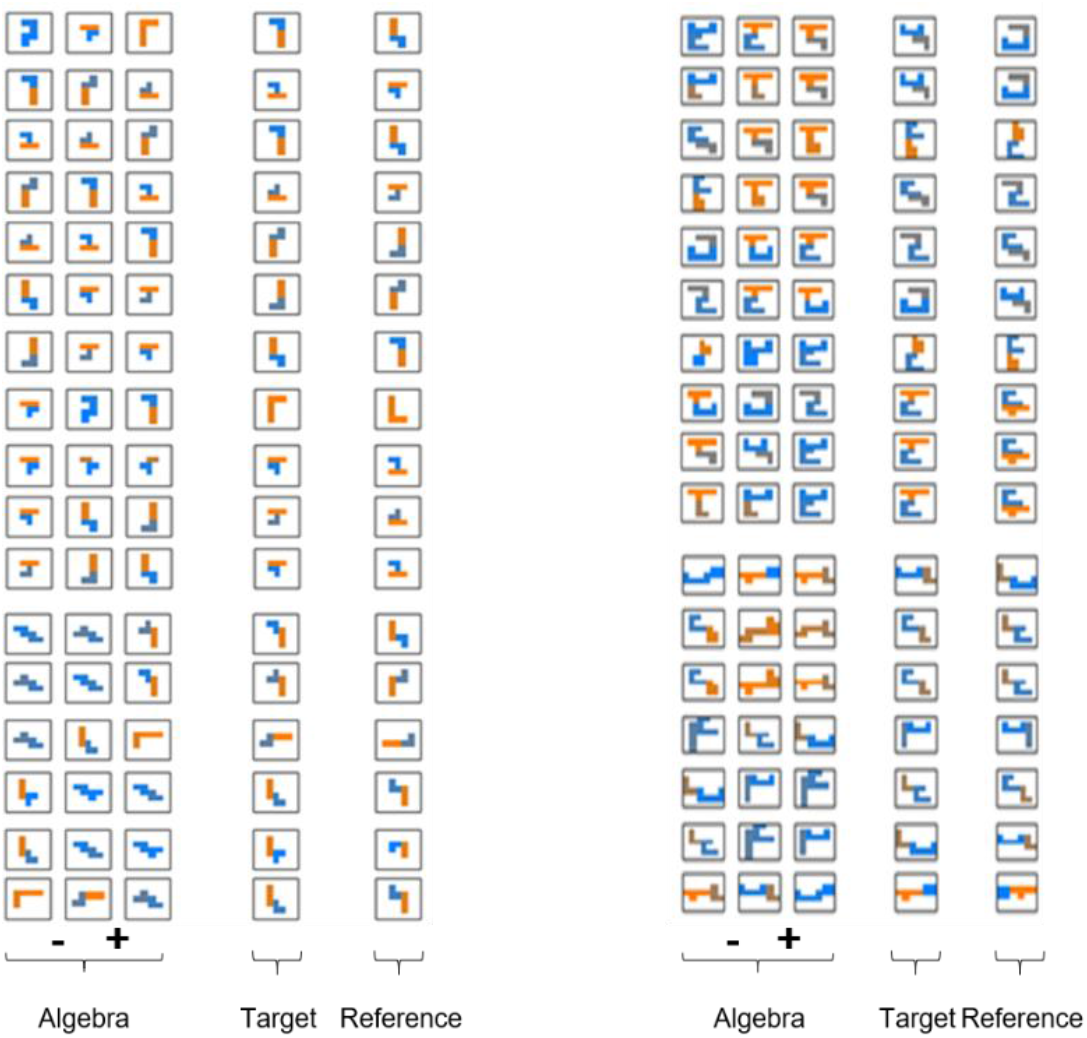
All non-hierarchical (left) and hierarchical (right) ‘silhouette algebra’ trials.

**Supplementary Figure 4.**
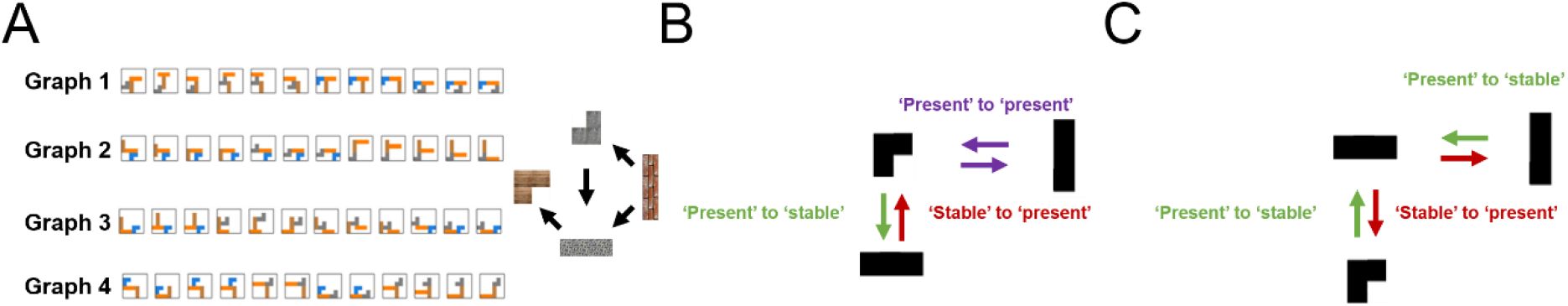
(A) Different stimulus sets used in the MEG task. Subjects were randomly assigned to one of these stimulus sets. (B) In half of the trials, the stable building block was not in the middle of the silhouette. (C) In the other half of trials, the stable building block was in the middle, such that there was no ‘present to present’ building block connection.

**Supplementary Figure 5.**
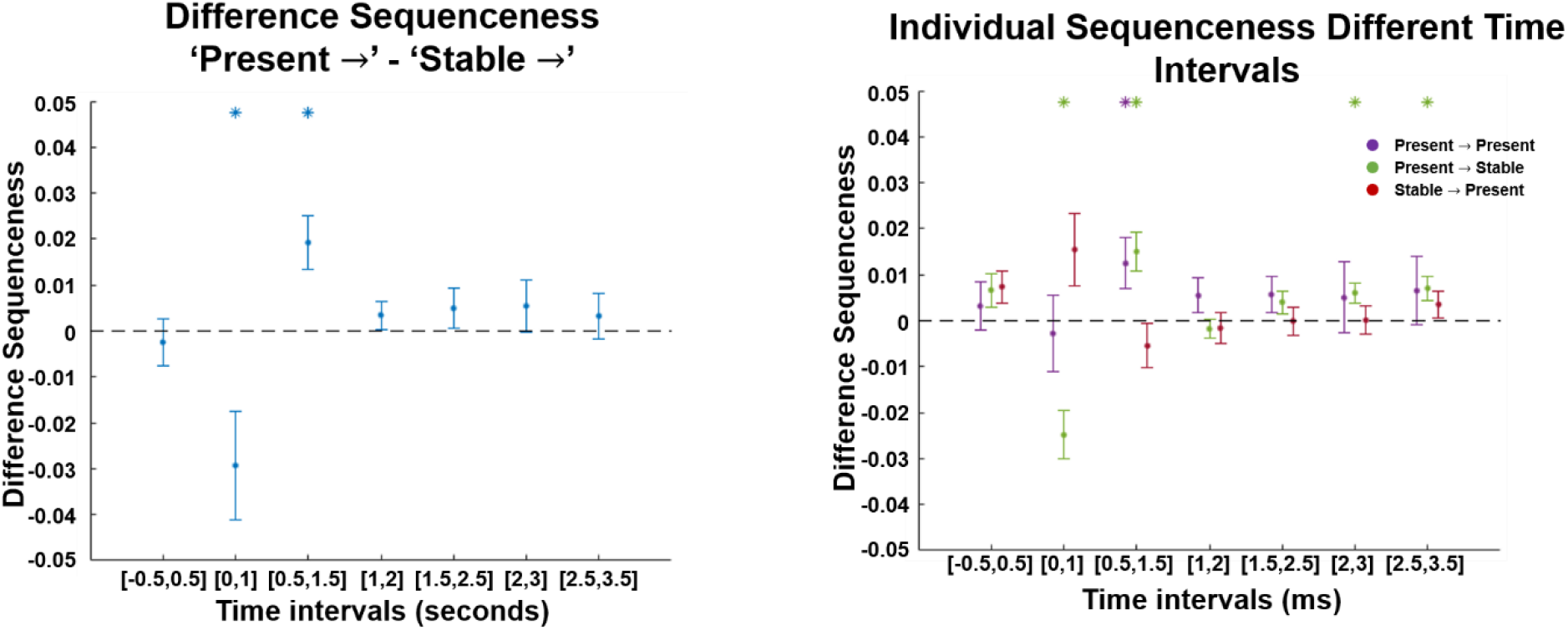
We investigated the difference between sequences starting either from the stable or the present building blocks for different time-intervals of the planning period, and found an early predominance of replay starting from the ‘stable’ building block (0 – 1000ms) followed by a predominance of replay starting from the ‘present’ building blocks (500ms-1500ms) during planning (left). Assessing the individual contributions of the different types of neural replay to these differences, we found a marked decrease of sequences toward the ‘stable’ building block early during planning (0-1000ms) followed by a predominance of sequences starting from the ‘present’ building blocks (500-1500ms). We also found a specific predominance of sequences from ‘present’ to the ‘stable’ building block during intervals at the end of the planning period (2000-3000ms and 2500-3500ms) before subjects entered the decision phase of the task.

## References

Aly, M., Ranganath, C., & Yonelinas, A. P. (2013). Detecting changes in scenes: The hippocampus is critical for strength-based perception. Neuron, 78(6), 1127–1137. https://doi.org/10.1016/j.neuron.2013.04.018

Ambrose, R. E., Pfeiffer, B. E., & Foster, D. J. (2016). Reverse Replay of Hippocampal Place Cells Is Uniquely Modulated by Changing Reward. Neuron, 91(5), 1124–1136. https://doi.org/10.1016/j.neuron.2016.07.047

Antonov, G. K., Gagne, C., Eldar, E., & Dayan, P. (2021). Optimism and Pessimism in Optimised Replay. BioRxiv, 2021.04.27.441454. https://doi.org/10.1101/2021.04.27.441454

Bapst, V., Sanchez-Gonzalez, A., Doersch, C., Stachenfeld, K. L., Kohli, P., Battaglia, P. W., & Hamrick, J. B. (2019). Structured agents for physical construction. ArXiv:1904.03177 [Cs]. http://arxiv.org/abs/1904.03177

Baram, A. B., Muller, T. H., Nili, H., Garvert, M. M., & Behrens, T. E. J. (2020). Entorhinal and ventromedial prefrontal cortices abstract and generalize the structure of reinforcement learning problems. Neuron. https://doi.org/10.1016/j.neuron.2020.11.024

Barron, H. C., Dolan, R. J., & Behrens, T. E. J. (2013). Online evaluation of novel choices by simultaneous representation of multiple memories. Nature Neuroscience, 16(10), 1492– 1498. https://doi.org/10.1038/nn.3515

Barron, H. C., Garvert, M. M., & Behrens, T. E. J. (2016). Repetition suppression: A means to index neural representations using BOLD? Philosophical Transactions of the Royal Society of London. Series B, Biological Sciences, 371(1705). https://doi.org/10.1098/rstb.2015.0355

Barry, D. N., & Love, B. C. (2021). A neural network account of memory replay and knowledge consolidation. BioRxiv, 2021.05.25.445587. https://doi.org/10.1101/2021.05.25.445587

Battaglia, P. W., Hamrick, J. B., Bapst, V., Sanchez-Gonzalez, A., Zambaldi, V., Malinowski, M., Tacchetti, A., Raposo, D., Santoro, A., Faulkner, R., Gulcehre, C., Song, F., Ballard, A., Gilmer, J., Dahl, G., Vaswani, A., Allen, K., Nash, C., Langston, V., … Pascanu, R. (2018). Relational inductive biases, deep learning, and graph networks. ArXiv:1806.01261 [Cs, Stat]. http://arxiv.org/abs/1806.01261

Behrens, T. E. J., Muller, T. H., Whittington, J. C. R., Mark, S., Baram, A. B., Stachenfeld, K. L., & Kurth-nelson, Z. (2018). What Is a Cognitive Map? Organizing Knowledge for Flexible Behavior. Neuron, 100(2), 490–509. https://doi.org/10.1016/j.neuron.2018.10.002

Bongioanni, A., Folloni, D., Verhagen, L., Sallet, J., Klein-Flügge, M. C., & Rushworth, M. F. S. (2021). Activation and disruption of a neural mechanism for novel choice in monkeys. Nature, 1–5. https://doi.org/10.1038/s41586-020-03115-5

Brunec, I. K., & Momennejad, I. (2021). Predictive Representations in Hippocampal and Prefrontal Hierarchies. BioRxiv, 786434. https://doi.org/10.1101/786434

Buckner, R. L., Goodman, J., Burock, M., Rotte, M., Koutstaal, W., Schacter, D., Rosen, B., & Dale, A. M. (1998). Functional-anatomic correlates of object priming in humans revealed by rapid presentation event-related fMRI. Neuron, 20(2), 285–296. https://doi.org/10.1016/s0896-6273(00)80456-0

Buzsáki, G. (2015). Hippocampal sharp wave-ripple: A cognitive biomarker for episodic memory and planning. Hippocampus, 25(10), 1073–1188. https://doi.org/10.1002/hipo.22488

Constantinescu, A. O., O’Reilly, J. X., & Behrens, T. E. J. (2016). Organizing conceptual knowledge in humans with a gridlike code. Science (New York, N.Y.), 352(6292), 1464–1468. https://doi.org/10.1126/science.aaf0941

Córdova, N. I., Turk-Browne, N. B., & Aly, M. (2019). Focusing on what matters: Modulation of the human hippocampus by relational attention. Hippocampus, 29(11), 1025–1037. https://doi.org/10.1002/hipo.23082

Dayan, P., Hinton, G. E., Neal, R. M., & Zemel, R. S. (1995). The Helmholtz machine. Neural Computation, 7(5), 889–904.

Diba, K., & Buzsáki, G. (2007a). Forward and reverse hippocampal place-cell sequences during ripples. Nature Neuroscience, 10(10), 1241–1242. https://doi.org/10.1038/nn1961

Diba, K., & Buzsáki, G. (2007b). Forward and reverse hippocampal place-cell sequences during ripples. Nature Neuroscience, 10(10), 1241–1242. https://doi.org/10.1038/nn1961

Eickhoff, S. B., Paus, T., Caspers, S., Grosbras, M.-H., Evans, A. C., Zilles, K., & Amunts, K. (2007). Assignment of functional activations to probabilistic cytoarchitectonic areas revisited. NeuroImage, 36(3), 511–521. https://doi.org/10.1016/j.neuroimage.2007.03.060

Eickhoff, S. B., Stephan, K. E., Mohlberg, H., Grefkes, C., Fink, G. R., Amunts, K., & Zilles, K. (2005). A new SPM toolbox for combining probabilistic cytoarchitectonic maps and functional imaging data. NeuroImage, 25(4), 1325–1335. https://doi.org/10.1016/j.neuroimage.2004.12.034

Eldar, E., Bae, G. J., Kurth-Nelson, Z., Dayan, P., & Dolan, R. J. (2018). Magnetoencephalography decoding reveals structural differences within integrative decision processes. Nature Human Behaviour, 2(9), 670–681. https://doi.org/10.1038/s41562-018-0423-3

Eslami, S. M. A., Jimenez Rezende, D., Besse, F., Viola, F., Morcos, A. S., Garnelo, M., Ruderman, A., Rusu, A. A., Danihelka, I., Gregor, K., Reichert, D. P., Buesing, L., Weber, T., Vinyals, O., Rosenbaum, D., Rabinowitz, N., King, H., Hillier, C., Botvinick, M., … Hassabis, D. (2018). Neural scene representation and rendering. Science (New York, N.Y.), 360(6394), 1204–1210. https://doi.org/10.1126/science.aar6170

Evans, T., & Burgess, N. (2020). Replay as structural inference in the hippocampal-entorhinal system. BioRxiv.

Finke, R. A., & Slayton, K. (1988). Explorations of creative visual synthesis in mental imagery. Memory & Cognition, 16(3), 252–257. https://doi.org/10.3758/BF03197758

Foster, D. J. (2017). Replay Comes of Age. Annual Review of Neuroscience, 40(1), 581–602. https://doi.org/10.1146/annurev-neuro-072116-031538

Foster, D. J., & Wilson, M. A. (2006). Reverse replay of behavioural sequences in hippocampal place cells during the awake state. Nature, 440(7084), 680–683. https://doi.org/10.1038/nature04587

Friston, K., & Buzsáki, G. (2016). The Functional Anatomy of Time: What and When in the Brain. Trends in Cognitive Sciences, 20(7), 500–511. https://doi.org/10.1016/j.tics.2016.05.001

Garvert, M. M., Dolan, R. J., & Behrens, T. E. J. (2017). A map of abstract relational knowledge in the human hippocampal–entorhinal cortex. ELife, 6, 1–20. https://doi.org/10.7554/eLife.17086

Gupta, A. S., van der Meer, M. A. A., Touretzky, D. S., & Redish, A. D. (2010). Hippocampal replay is not a simple function of experience. Neuron, 65(5), 695–705. https://doi.org/10.1016/j.neuron.2010.01.034

Hamrick, J. B., Allen, K. R., Bapst, V., Zhu, T., McKee, K. R., Tenenbaum, J. B., & Battaglia, P. W. (2018). Relational inductive bias for physical construction in humans and machines. ArXiv:1806.01203 [Cs, Stat]. http://arxiv.org/abs/1806.01203

Hartley, T., Lever, C., Burgess, N., & O ‘keefe, J. (2014). Space in the brain: How the hippocampal formation supports spatial cognition. Philosophical Transactions of the Royal Society of London, 369(1635), 20120510–20120510. https://doi.org/10.1098/rstb.2012.0510

Hassabis, D., Kumaran, D., & Maguire, E. A. (2007). Using imagination to understand the neural basis of episodic memory. The Journal of Neuroscience: The Official Journal of the Society for Neuroscience, 27(52), 14365–14374. https://doi.org/10.1523/JNEUROSCI.4549-07.2007

Hassabis, D., & Maguire, E. A. (2009). The construction system of the brain. Philosophical Transactions of the Royal Society of London. Series B, Biological Sciences, 364(1521), 1263– 1271. https://doi.org/10.1098/rstb.2008.0296

Hunt, L. T., Malalasekera, W. M. N., de Berker, A. O., Miranda, B., Farmer, S. F., Behrens, T. E. J., & Kennerley, S. W. (2018). Triple dissociation of attention and decision computations across prefrontal cortex. Nature Neuroscience, 21(10), 1471–1481. https://doi.org/10.1038/s41593-018-0239-5

Johnson, A., & Redish, A. D. (2007). Neural Ensembles in CA3 Transiently Encode Paths Forward of the Animal at a Decision Point. Journal of Neuroscience, 27(45), 12176–12189. https://doi.org/10.1523/JNEUROSCI.3761-07.2007

Kay, K., Chung, J. E., Sosa, M., Schor, J. S., Karlsson, M. P., Larkin, M. C., Liu, D. F., & Frank, L. M. (2020). Constant Sub-second Cycling between Representations of Possible Futures in the Hippocampus. Cell, 180(3), 552-567.e25. https://doi.org/10.1016/j.cell.2020.01.014

Kemp, C., & Tenenbaum, J. B. (2008). The discovery of structural form. Proceedings of the National Academy of Sciences of the United States of America, 105(31), 10687–10692. https://doi.org/10.1073/pnas.0802631105

Kriegeskorte, N., Goebel, R., & Bandettini, P. (2006). Information-based functional brain mapping. Proceedings of the National Academy of Sciences, 103(10), 3863–3868. https://doi.org/10.1073/pnas.0600244103

Kriegeskorte, N., Mur, M., & Bandettini, P. (2008). Representational similarity analysis—Connecting the branches of systems neuroscience. Frontiers in Systems Neuroscience, 2, 4–4. https://doi.org/10.3389/neuro.06.004.2008

Kurth-Nelson, Z., Economides, M., Dolan, R. J., & Dayan, P. (2016). Fast Sequences of Non-spatial State Representations in Humans. Neuron, 91(1), 194–204. https://doi.org/10.1016/j.neuron.2016.05.028

Lake, B. M. (2019). Compositional generalization through meta sequence-to-sequence learning. ArXiv:1906.05381 [Cs]. http://arxiv.org/abs/1906.05381

Lake, B., Salakhutdinov, R., & Tenenbaum, J. (2015). Human-level concept learning through probabilistic program induction. Science (New York, N.Y.), 350(6266), 1332–1338. https://doi.org/10.1126/science.aab3050

Lee, A. C. H., Buckley, M. J., Pegman, S. J., Spiers, H., Scahill, V. L., Gaffan, D., Bussey, T. J., Davies, R. R., Kapur, N., Hodges, J. R., & Graham, K. S. (2005). Specialization in the medial temporal lobe for processing of objects and scenes. Hippocampus, 15(6), 782–797. https://doi.org/10.1002/hipo.20101

Lee, A. C. H., Bussey, T. J., Murray, E. A., Saksida, L. M., Epstein, R. A., Kapur, N., Hodges, J. R., & Graham, K. S. (2005). Perceptual deficits in amnesia: Challenging the medial temporal lobe ‘mnemonic’ view. Neuropsychologia, 43(1), 1–11. https://doi.org/10.1016/j.neuropsychologia.2004.07.017

Liu, Y., Dolan, R. J., Kurth-Nelson, Z., & Behrens, T. E. (2019). Human replay spontaneously reorganizes experience. Cell, 178(3), 640–652.

Liu, Y., Dolan, R. J., Penagos-Vargas, H. L., Kurth-Nelson, Z., & Behrens, T. (2020). Measuring Sequences of Representations with Temporally Delayed Linear Modelling. BioRxiv, 2020.04.30.066407. https://doi.org/10.1101/2020.04.30.066407

Liu, Y., Mattar, M. G., Behrens, T. E. J., Daw, N. D., & Dolan, R. J. (2021). Experience replay is associated with efficient nonlocal learning. Science, 372(6544). https://doi.org/10.1126/science.abf1357

Luyckx, F., Nili, H., Spitzer, B., & Summerfield, C. (2019). Neural structure mapping in human probabilistic reward learning. ELife, 8, e42816. https://doi.org/10.7554/eLife.42816

Manns, J. R., & Eichenbaum, H. (2006). Evolution of declarative memory. Hippocampus, 16(9), 795– 808. https://doi.org/10.1002/hipo.20205

Mark, S., Moran, R., Parr, T., Kennerley, S. W., & Behrens, T. E. J. (2020). Transferring structural knowledge across cognitive maps in humans and models. Nature Communications, 11(1), 4783. https://doi.org/10.1038/s41467-020-18254-6

Mattar, M. G., & Daw, N. D. (2018). Prioritized memory access explains planning and hippocampal replay. Nature Neuroscience, 21(11), 1609–1617.

Miller, K. J., Botvinick, M. M., & Brody, C. D. (2017). Dorsal hippocampus contributes to model-based planning. Nature Neuroscience, 20(9), 1269–1276. https://doi.org/10.1038/nn.4613

Morton, N. W., Schlichting, M. L., & Preston, A. R. (2020). Representations of common event structure in medial temporal lobe and frontoparietal cortex support efficient inference. Proceedings of the National Academy of Sciences, 117(47), 29338–29345. https://doi.org/10.1073/pnas.1912338117

Nichols, T. E., & Holmes, A. P. (2002). Nonparametric permutation tests for functional neuroimaging: A primer with examples. Human Brain Mapping, 15(1), 1–25. https://doi.org/10.1002/hbm.1058

Nili, H., Wingfield, C., Walther, A., Su, L., Marslen-Wilson, W., & Kriegeskorte, N. (2014). A Toolbox for Representational Similarity Analysis. PLOS Computational Biology, 10(4), e1003553. https://doi.org/10.1371/journal.pcbi.1003553

O’keefe, J., & Nadel, L. (1978). The hippocampus as a cognitive map. Oxford: Clarendon Press.

Ólafsdóttir, H. F., Barry, C., Saleem, A. B., Hassabis, D., & Spiers, H. J. (2015). Hippocampal place cells construct reward related sequences through unexplored space. ELife, 4, e06063. https://doi.org/10.7554/eLife.06063

Orbán, G., Fiser, J., Aslin, R. N., & Lengyel, M. (2008). Bayesian learning of visual chunks by human observers. Proceedings of the National Academy of Sciences, 105(7), 2745–2750. https://doi.org/10.1073/PNAS.0708424105

Pfeiffer, B. E., & Foster, D. J. (2013). Hippocampal place-cell sequences depict future paths to remembered goals. Nature, 497(7447), 74–79. https://doi.org/10.1038/nature12112

Ruiz, N. A., Meager, M. R., Agarwal, S., & Aly, M. (2020). The Medial Temporal Lobe Is Critical for Spatial Relational Perception. Journal of Cognitive Neuroscience, 32(9), 1780–1795. https://doi.org/10.1162/jocn_a_01583

Schuck, N. W., & Niv, Y. (2019). Sequential replay of nonspatial task states in the human hippocampus. Science (New York, N.Y.), 364(6447). https://doi.org/10.1126/science.aaw5181

Stachenfeld, K. L., Botvinick, M. M., & Gershman, S. J. (2017). The hippocampus as a predictive map. Nature Neuroscience, 20(11), 1643.

Sutton, R. S., & Barto, A. G. (1998). Reinforcement Learning: An Introduction. Advances in Cancer Research, 104, 322–322. https://doi.org/10.1016/S0065-230X(09)04001-9

Tolman, E. C. (1948). Cognitive maps in rats and men. Psychological Review, 55(4), 189–208. https://doi.org/10.1037/h0061626

Ullman, T. D., Spelke, E., Battaglia, P., & Tenenbaum, J. B. (2017). Mind Games: Game Engines as an Architecture for Intuitive Physics. Trends in Cognitive Sciences, 21(9), 649–665. https://doi.org/10.1016/j.tics.2017.05.012

Vikbladh, O. M., Meager, M. R., King, J., Blackmon, K., Devinsky, O., Shohamy, D., Burgess, N., & Daw, N. D. (2019). Hippocampal Contributions to Model-Based Planning and Spatial Memory. Neuron, 102(3), 683-693.e4. https://doi.org/10.1016/j.neuron.2019.02.014

Whittington, J. C. R., Muller, T. H., Mark, S., Chen, G., Barry, C., Burgess, N., & Behrens, T. E. J. (2020). The Tolman-Eichenbaum Machine: Unifying Space and Relational Memory through Generalization in the Hippocampal Formation. Cell, 183(5), 1249-1263.e23. https://doi.org/10.1016/j.cell.2020.10.024

Winkler, A. M., Ridgway, G. R., Webster, M. A., Smith, S. M., & Nichols, T. E. (2014). Permutation inference for the general linear model. NeuroImage, 92, 381–397. https://doi.org/10.1016/j.neuroimage.2014.01.060

Yamins, D. L. K., & DiCarlo, J. J. (2016). Using goal-driven deep learning models to understand sensory cortex. Nature Neuroscience, 19(3), 356–365. https://doi.org/10.1038/nn.4244

